# Adult neurogenesis promotes efficient, nonspecific search strategies in a spatial alternation water maze task

**DOI:** 10.1101/637462

**Authors:** Ru Qi Yu, Matthew Cooke, Jiaying Zhao, Jason S. Snyder

**Affiliations:** Department of Psychology, Djavad Mowafaghian Centre for Brain Health, University of British Columbia, 2211 Wesbrook Mall, Vancouver, British Columbia, Canada V6T 2B5

**Keywords:** spatial learning, regularities, navigation, strategy, hippocampus

## Abstract

Goal-directed navigation requires learning strategies that are efficient and minimize costs. In some cases it may be desirable to flexibly adjust behavioral responses depending on the cues that vary from one episode to the next. In others, successful navigation might be achieved with inflexible, habit-like responses that reduce cognitive load. Adult neurogenesis is believed to contribute to the spatial processing functions of the hippocampus, particularly when behavioral flexibility is required. However, little is known about the role of neurogenesis in spatial navigation when goals are unpredictable or change according to certain rules. We hypothesized that neurogenesis is necessary in a spatial navigation task that involves different patterns of reinforcement. Intact and neurogenesis-deficient rats were trained to escape to one of two possible platform locations in a spatial water maze. The platform either repeated in the same location for all trials in a day, alternated between two locations across trials, or randomly moved between the two locations. Neurogenesis selectively enhanced escape performance in the alternating condition, but not by improving platform choice accuracy. Instead, neurogenesis-intact rats made fewer search errors and developed an efficient habit-like strategy where they consistently swam to a preferred location. If the platform was not present, they proceeded to the other possible location. In contrast, neurogenesis-deficient rats were indecisive and navigationally less-efficient. Thus, in conditions where goals follow a predictable spatiotemporal pattern, adult neurogenesis promotes the adoption of navigation strategies that are spatially nonspecific but, nonetheless, accurate and efficient.

## 1. Introduction

Adult neurogenesis has been widely studied in the context of hippocampal functions in memory. In spatial navigation tasks such as the water maze and the active place avoidance task, neurogenesis is typically not required for rodents to learn a single spatial location [1–4]. Instead, neurogenesis is often more important for spatial reversal, i.e. ignoring a location that no longer offers escape and learning to approach a new escape location [5–7]. A similar role in spatial reversal has been observed in an appetitive operant touchscreen task, where the rewarded spatial choice changes across blocks of trials [8]. This pattern of flexible behavior is also apparent in studies showing that neurogenesis is important for discriminative fear conditioning [9–11] and learning lists of odors that have conflicting reward associations [12]. In all of these tasks there is a high degree of similarity between experiences, making it difficult to choose between competing options.

Natural situations can be even more complex, because we are often repeatedly faced with competing choices that are also imperfectly associated with outcomes. With respect to spatial navigation, the optimal or most efficient response may therefore entail more than simply forming an accurate memory, or performing a simple reversal. For example, efficiently finding your car in a parking garage requires a strategy for distinguishing multiple competing goals, if you park in different locations each day. When situations repeat themselves, however, there may be temporal regularities that can be exploited to learn the appropriate response or develop an optimal strategy. Indeed, the hippocampus is critical for remembering the spatiotemporal order of events and guiding navigational behavior based on memory for past experiences [13–15].

Despite the role of neurogenesis in flexible spatial behavior, and forming associations over time, it is unknown whether neurogenesis contributes to flexible navigation when spatial goals change according to a temporal pattern. Additionally, it is unknown whether adult neurogenesis modifies navigational strategies when spatial goals vary in their degree of predictability. To examine whether adult neurogenesis promotes learning of spatiotemporal patterns we subjected neurogenesis-deficient GFAP-TK rats to a water maze task where the platform moved between two locations with minimal temporal interference (repeats in the same location each day), higher interference but still predictable (alternates between two locations across trials), and a random temporal pattern that could not be learned. We hypothesized that neurogenesis would be specifically required for learning the alternating pattern since delayed spatial alternation and nonmatch behaviors are hippocampal dependent [16,17]. Moreover, the alternating pattern is essentially a series of delayed non-match to place trials, which have been suggested to require adult neurogenesis, though choice behaviors for that study were not reported [18]. While neurogenesis-intact rats escaped faster in the alternating condition, a detailed analysis of their spatial search patterns and decision-making behaviors revealed that they did not in fact alternate. Instead they were generally more accurate in their platform approaches and they targeted the two platform locations in an identical sequence on each trial, which enabled them to rapidly escape regardless of where the platform was located. These results suggest that adult neurogenesis promotes efficient search strategies that minimize cognitive effort.

## 2. Methods

### 2.1 Subjects and treatment

All procedures were approved by the Animal Care Committee at the University of British Columbia and conducted in accordance with the Canadian Council on Animal Care guidelines regarding humane and ethical treatment of animals. Experimental Long Evans rats were generated and maintained in the Department of Psychology animal facility with a 12-hr light/dark schedule and lights on at 6:00 a.m. Breeding occurred in large polyurethane cages (47 cm × 37 cm × 21 cm) containing a polycarbonate tube, aspen chip bedding, and ad libitum rat chow and water. Litters ranged from 8 to 18 pups and pups from different litters were distributed equally among experimental groups. Both male and female breeders remained with the litters until P21, when offspring were weaned to 2 per cage in smaller polyurethane cages (25 cm × 37 cm × 21 cm).

Forty-nine male rats were used in this study, consisting of 26 transgenic GFAP-TK rats (TK, neurogenesis-intact) and 23 wild-type (WT, neurogenesis-deficient) littermates. In these rats, oral treatment with valganciclovir suppresses adult neurogenesis specifically in TK rats but leaves WT rats intact [19]. At 8 weeks of age, all animals received drug treatment twice per week for 8 weeks. For each treatment, rats were given 7.5 mg of valganciclovir in a 0.5 g pellet consisting of a 1:1 mixture of ground chow and peanut butter. The treatment was terminated before the first day of behavioral testing.

### 2.2 General behavioral testing procedures

WT and TK rats were trained on variants of the spatial water maze where the escape platform was located in 1 of 2 locations according to a repeating, alternating, or random pattern. The water maze was a circular pool, 2 meters in diameter, located in a dimly-lit testing room approximately 4 meters × 6 meters in size. Large spatial cues, distal to the maze, were present throughout the room (symbols on the walls, a door, the computer, cupboards). The pool water was room temperature (~22°C) and made opaque with white tempera paint. The pool was divided into 4 equal-sized invisible quadrants (NW,NE,SE,SW) and a hidden white escape platform could appear in the center of a given quadrant (circular platform, 10 cm diameter, submerged 5 cm). For all testing conditions, rats were given 10 trials per day for 15 days. On each trial, the platform could appear in only 1 of 2 possible locations, which were pre-determined for each rat and counter-balanced across conditions (Fig. 1). For half of the rats in each condition the two possible platform locations were in the same half of the pool (e.g. NW and SW). In this case the rat was released from the opposite side of the pool, equidistant from both locations (e.g. E). For the other half of the rats, the two possible platform locations were in diagonally opposite quadrants (e.g. SW and NE). In this case the rat was also released equidistant from the two possible platform locations (e.g. NW or SE). For a given rat, platform locations and release points were constant for all 150 trials of the experiment.

At the beginning of each testing day the 2 rats in a cage were brought to the testing room and put into separate holding cages. Testing trials alternated between the 2 cage mates (i.e. rat 1 trial 1, rat 2 trial 1, rat 1 trial 2…) and it took ~20 s to set up the platform position in between trials. A structural base was positioned in each of the potential platform locations and the escape platform was positioned by attaching to the top of the relevant base. In this way, rats could touch the base of the alternate potential platform location but they could not mount it for escape purposes. By days 10-15 of training (when most analyses were conducted), the intertrial interval was 76 seconds on average. On each trial, the rat was released facing the pool wall and the trial ended when the rat stayed on the platform for 0.5 s, or when 70 s had elapsed (at which point the rat was guided to the platform). Rats were retrieved after spending 20 s on the platform, towel dried and placed into their holding cage. After the days’ trials were complete, the cagemates were put back together into their home cage and returned to the animal colony. The genotype of two rats in a cage was not predetermined. Instead, rats were weaned prior to genotyping resulting in a random distribution of WT and TK rats across cages. Generally, cagemates were assigned to the same testing condition except for three cages where each rat in the cage was assigned to a different condition for counter-balancing purposes.

### 2.3 Spatial reinforcement patterns

As mentioned above, the platform was located in only 1 of 2 possible locations for each rat. In the *repeating* condition, the platform appeared in the same location for all 10 trials in a given day, and on the next day the platform appeared in the other location for all trials. In the *alternating* condition, the platform location perfectly alternated between the two possible locations across trials in a day. In the *random* condition, the platform appeared randomly in the two possible locations each day, with the constraint that both platform locations were equally reinforced within a day.

Rats in the 3 conditions were therefore treated equally in terms of the number of potential platform locations, the spatial locations of the platforms, release points within the pool, and the time of testing. The primary difference among the conditions was the temporal pattern by which the 2 possible platform locations were reinforced. The repeating pattern of reinforcement is the most predictable pattern, and likely the easiest to learn given that the pattern only changed across days. The alternating pattern is likely to lead to more interference due to the spatiotemporal switching between the 2 locations. However, we hypothesized that WT rats, but possibly not TK rats, would still perform well in the alternating condition. Finally, given the lack of predictability in the random condition, we expected WT and TK rats to perform poorly and fail to predict the correct location of the platform on a given trial.

**Figure 1:**
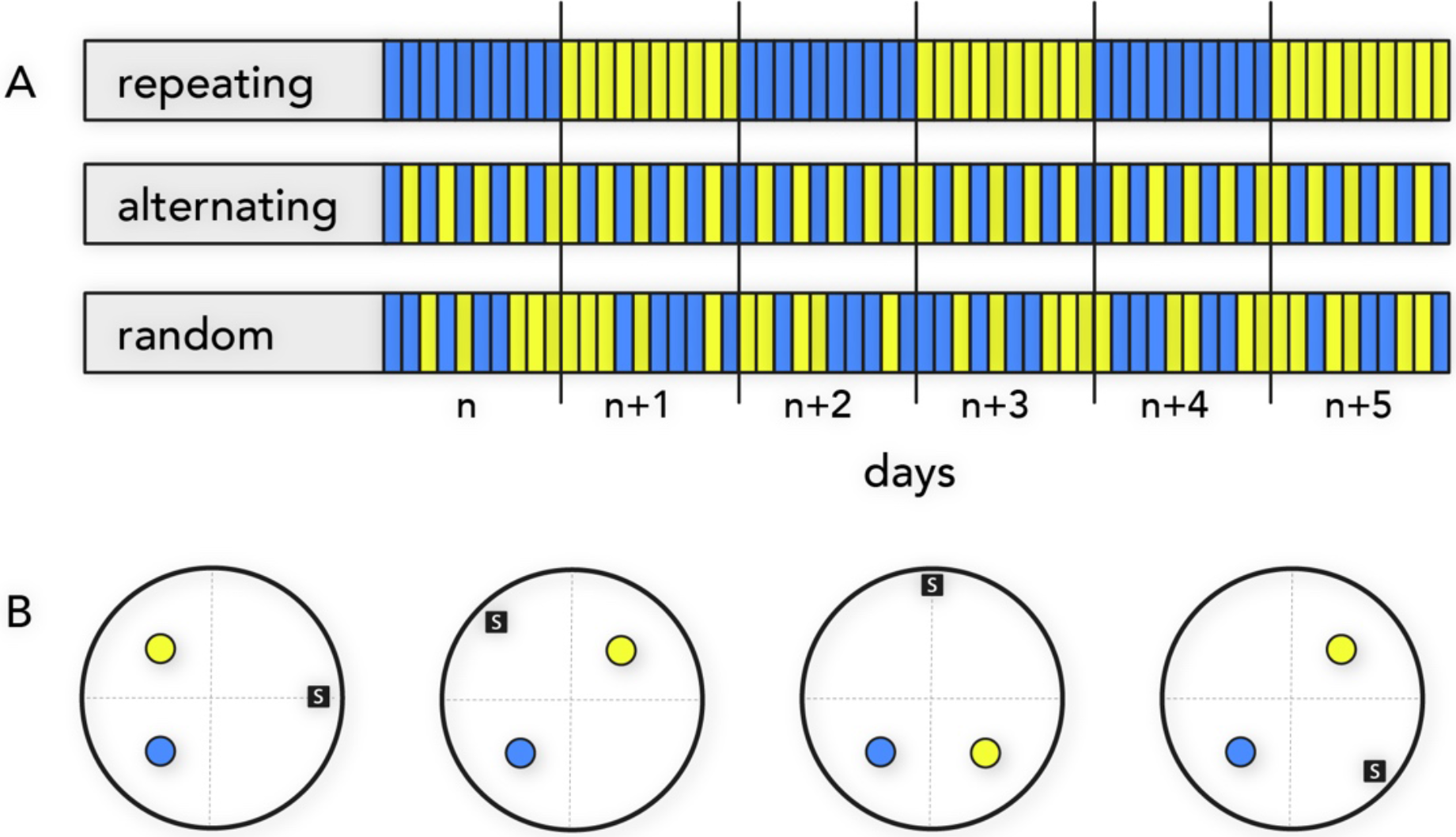
Behavioral paradigm. A) Schematic illustration of platform reinforcement patterns. All rats received 10 trials/day where the escape platform moved between 2 possible locations according to a repeating, alternating or random pattern. B) The four combinations of platform locations and start points used in all 3 conditions. Start location is indicated by “s”.

### 2.4 Spatial behavior analyses

Two dimensional swim paths were tracked from above at 25 Hz using EthoVision XT10 (Noldus). These data were used to calculate spatial and temporal performance metrics using native functions within Ethovision or with custom analyses of swim paths relative to pool geometry and landmarks. Standard measures included latency to reach the platform, path length, swim speed, proximity (average distance) to the platform locations, heading angle, latency to reach 40 cm zones centered on the possible platform locations, and zone crossings [20,21].

Heading angle quantified the swimming direction, relative to a direct path between the rat and the platform, at 9 equally-spaced points in time, p, along the swim path (corresponding to 0%, 11%, 22% … 100% of the trial duration). At a given point in the trajectory, p, the heading angle was the angular difference between a line connecting p to the platform and a line connecting p to the next point in the trajectory, p+1. The heading angle therefore provides a measure of navigational accuracy over the course of each swim [22].

Analyses of swim paths were conducted to assess the degree of spatial search specificity [5,23,24]. We focussed our analyses on days 10-15 of testing because animals were well-trained by then and did not display search strategies associated with learning procedural aspects of the task (e.g thigmotaxis). In the analysis, we focused on strategies that revealed the degree of search specificity for the correct platform. The first strategy was direct swim, defined as trials where the average heading angle error relative to the platform was ≤ 35° and the cumulative search error was ≤ 125 [21,25]. Focal search was defined as a swim path where centroid (mean of all the data points) was within 31.5 cm from the platform (i.e. relatively accurate) and the average distance from the centroid is within 40.5 cm (i.e. focused). However, since animals were well trained, focal search was relatively rare and appeared as only a minor deviation from a direct path. These trials were therefore considered as direct swim, which collectively reflected near-perfect trajectories to the platform. The second strategy was indirect search, defined as trials that did not meet the criteria for direct swim but where rats displayed a cumulative search error of ≤ 350. These criteria typically detected swim trajectories that contained a single incorrect turn (for example, when rats approached the incorrect platform location before re-routing). The third strategy was spatially non-specific search, defined as trials that did not meet the above criteria, which typically contained 2 or more turns, indicating multiple search errors within a trial.

### 2.5 Histology

Soon after finishing the last day of training, rats were sacrificed and their brains were extracted for histological verification that neurogenesis was selectively reduced in the TK rats. Sections containing dentate gyrus were immunostained goat anti-doublecortin (1:250; Santa Cruz, Dallas, TX, sc-8066) to visualize immature, adult-born cells as we have done previously [26].

### 2.6 Statistical analyses

Repeated-measures ANOVA, with Holm-Sidak post-hoc comparisons, were used to analyze WT and TK rats’ behavior over days and trials. One sample t-tests were used to compare platform preference to chance levels. Two-way repeated-measures ANOVA was used to investigate platform preference × genotype differences. In all cases, significance was set at P=0.05.

## 3. Results

### 3.1 Neurogenesis reduction

As expected, immature DCX^+^ cells were greatly reduced in the TK rats compared to WT rats. In a separate study, conducted at the same time and using the same valganciclovir treatment regimen, we quantified neurogenesis in WT and TK rats and found a 95% reduction of neurogenesis in the TK rats [26]. Since we observed a qualitatively similar reduction in immature DCX^+^ cells here, we did not explicitly quantify neurogenesis in the current study but conclude that neurogenesis reduction was highly effective in the TK rats.

### 3.2 Task acquisition

General acquisition of task demands was assessed over the full 15 days of training for all 3 conditions (Supplementary Fig. 1). The latency to escape to the hidden platform decreased over days and was similar between WT and TK rats in all three conditions. Since latency did not reveal where animals searched, we also examined proximity (mean distance) to the platform location, which also improved across days of training in WT and TK rats. Since rats were faced with a choice of two platform locations, we examined the proximity to the incorrect platform location over days. While WT and TK rats were similar in the repeating and random conditions, TK rats’ swim paths were closer to the incorrect location in the later stages of training in the alternating condition (Supplementary Fig. 1).

### 3.3 Task performance

In the water maze, rats first learned procedural aspects of the task (e.g. avoid pool walls) prior to forming a precise spatial map [23]. Indeed, differences in proximity to the incorrect location only emerged once animals were well trained. We therefore focused our detailed analyses on days 10-15, once basic task demands had been mastered and performance had stabilized.

We first examined escape latencies across the 10 trials within each day, averaged across the last 6 days of testing (Fig. 2a-e). In the repeating condition, there was no difference between WT and TK rats in average latencies. Both groups showed a steep decline over trials 1-3 followed by a plateau in performance, indicating rapid learning of the platform location on each day. In the alternating condition, WT rats escaped faster than TK rats. In the random condition, WT and TK rats were not statistically different. A within genotype analysis revealed that TK rats in the alternating condition was not statistically different from the TK rats in the random condition, whereas WT rats in the repeating and alternating conditions performed comparably, and better than the random condition (Fig. 2e-d). On trial 1, both TK and WT rats in the repeating condition were slower to escape than in alternating or random conditions, suggesting that they perseverated at the previous day’s platform location, or they may have utilized different strategies for rapidly locating the platform when no previous trial information was available. Shorter escape latencies in the WT alternating condition were unlikely to reflect motor/performance differences in WT vs TK rats since swim speeds were comparable (WT: 34.5 cm/s, TK: 32.6 cm/s; T_16_=1.1, P=0.3) and a similar pattern of results was obtained when path lengths were analyzed (Supplementary Fig. 2).

To investigate search relative to the correct and incorrect platform locations, we examined proximity to the platform locations across trials (Fig. 2f-k). WT and TK rats did not differ in their proximity to the correct platform location, but differences emerged with respect to their proximity to the incorrect location. In the repeating condition, TK rats swam in closer proximity to the incorrect platform location on trial 2 compared to WT rats, indicating perseveration at the previous day’s escape location. In the alternating condition, TK rats consistently swam in closer proximity to the incorrect location, suggesting an impaired ability to discriminate the correct and incorrect platform locations.

**Figure 2:**
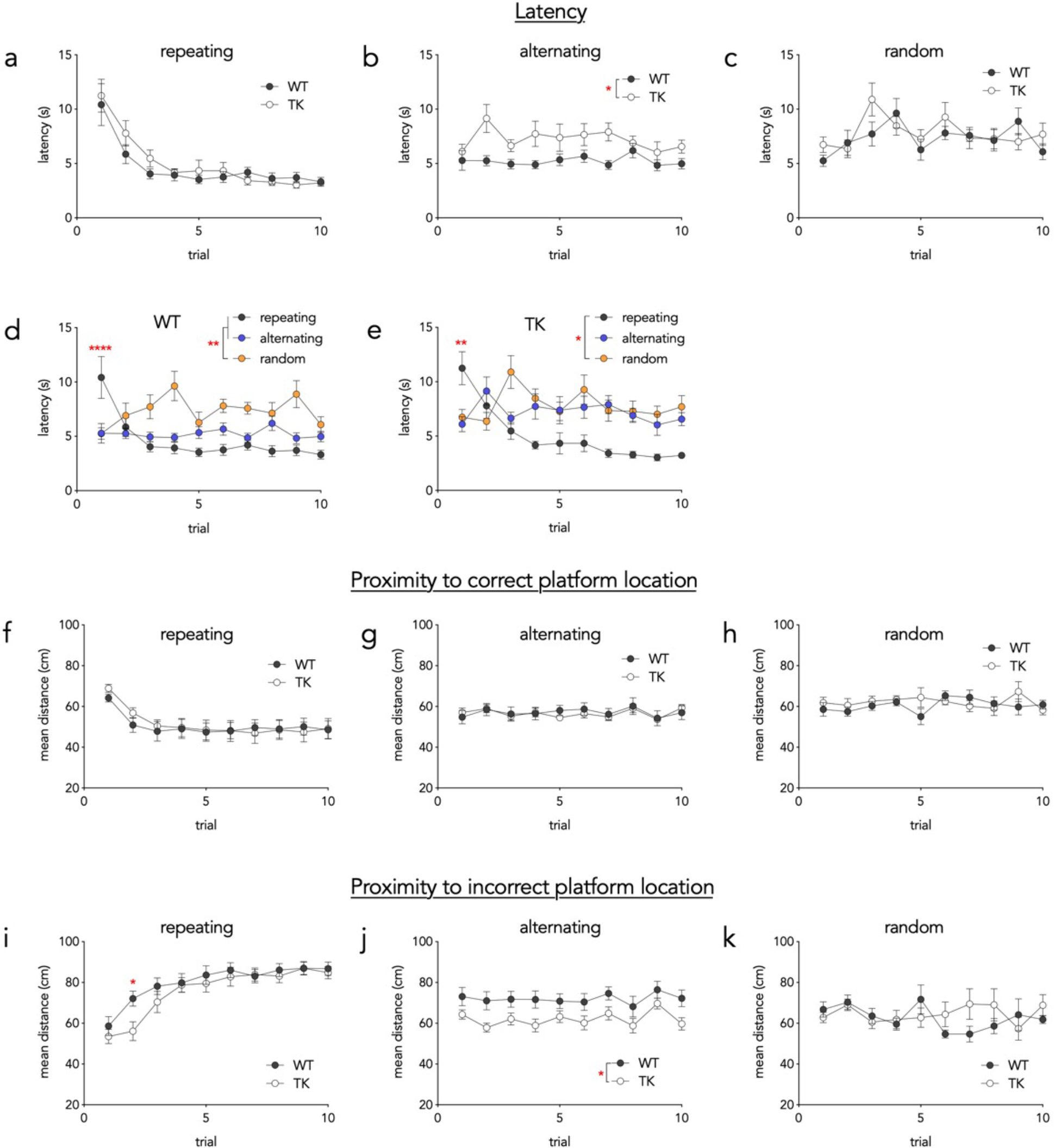
Performance on days 10-15. a) WT and TK rats in the repeating condition did not differ, and required 2 trials to reach asymptotic performance (effect of genotype, F_1,14_=0.4, P=0.55; effect of trial, F_9,126_=23, P<0.0001, trials 1 & 2 vs trials 3-10 all P<0.01). b) In the alternating condition, WT rats found the platform faster than TKs (F_1,16_=7, P=0.02). c) In the random condition, WT and TK rats found the platform equally fast (F_1,14_=0.3, P=0.6). d) WT rats in the random condition were generally slower to locate the platform that rats in the repeating and alternating conditions (effect of condition, F_2,21_=13, P=0.0002; **P<0.01 vs alternating and repeating groups). On trial 1, however, rats in the repeating condition were slower than both alternating and random groups (condition × trial interaction, F_18,189_=5.3, P<0.0001; post hoc comparisons both ****P<0.0001). e) TK rats in the repeating condition generally located the platform faster than rats in the random condition (effect of condition, F_2,23_=5.2, P=0.01; repeating vs random *P=0.02, repeating vs alternating P=0.05). On trial 1, TK rats in the repeating condition were slower to locate the platform (condition × trial interaction, F_18,207_=6.4, P<0.0001; post hoc comparisons both **P<0.01). On trials 4,6,7,8,10 rats in the repeating condition were faster to escape than rats in the alternating condition (all P<0.05). f) In the repeating condition, mean distance to the correct platform location decreased across trials (F_9,117_=19, P<0.0001) and was not difference between WT and TK rats (F_1,13_=0.03, P=0.9). g) In the alternating condition, WT and TK rats’ mean distance to the correct platform did not differ (F_1,16_=0.02, P=0.9). h) In the random condition, WT and TK rats’ mean distance to the correct platform did not differ (F_1,14_=0.3, P=0.6). i) In the repeating condition, mean distance to the incorrect platform location increased across trials (F_9,117_=37, P<0.0001) and was greater in WT than TK rats on trial 2 (trial × genotype interaction, F_9,117_=2.0, P=0.04; post hoc *P=0.04). g) In the alternating condition, WT rats’ mean distance to the incorrect platform was greater than TK rats (F_1,16_=6.1, P=0.02). h) In the random condition, WT and TK rats’ mean distance to the incorrect platform did not differ (genotype effect F_1,14_=0.2, P=0.6; trial × genotype interaction F_9,126_=0.04, post hoc WT vs TK all Ps>0.3).

### 3.4 Correct vs. incorrect platform choice

The results thus far suggest that adult neurogenesis is required for learning a simple spatial alternation task, consistent with previous evidence that the hippocampus is required for spatial alternation when there is a delay [16] (here, the average intertrial interval across all conditions, for days 10-15, was 76 seconds; for all conditions, WT vs TK: all Ps ≥ 0.11). If escape behavior was based on knowledge of the spatiotemporal pattern, rats should approach the correct platform location first on more than 50% of trials. We therefore quantified the proportion of trials where rats crossed a circular zone (40 cm diameter) that was centered on the correct location before crossing an equivalent zone centered on the incorrect location. We chose a zone that is larger than the platform because the front paws were several cm anterior to the portion of the body that was identified by the tracking software. A well-trained rat could therefore correctly reject the incorrect platform location but escape detection by the tracking software. We therefore used zones that extend ~1 body length from the edge of the platform locations to effectively capture platform approaches while avoiding false negatives. Analyses were limited to days 10-15 (to examine post-acquisition performance).

We first examined performance on trial 1 to determine whether initial choice was influenced by the platform location on the previous day (Fig. 3a). Consistent with the trial 1 latencies, path lengths and proximity data, rats in the repeating condition approached the correct location first on less than 50% of trials, indicating initial perseveration at the previous day’s location. Rats in the alternating condition chose the correct location on trial 1 at chance levels, even though the location was identical to trial 10 on the previous day. Rats in the random condition also approached the correct platform location at chance levels.

To determine if rats could learn and apply knowledge of spatiotemporal patterns once they know the day’s sequence we examined correct platform choice on trials 2-10 (Fig. 3b). Rats in the repeating condition swam to the correct location first on ~90% of trials, indicating successful reversal and rapid learning of the current location. In contrast, rats in the alternating and random conditions chose the correct location first at chance levels. There was no difference between WT and TK rats in any of the 3 conditions. These data indicate that only rats in the repeating condition were specifically targeting the correct platform and that faster escape latencies in alternating WT rats is not due to accurate choice (i.e. successful spatial alternation).

**Figure 3:**
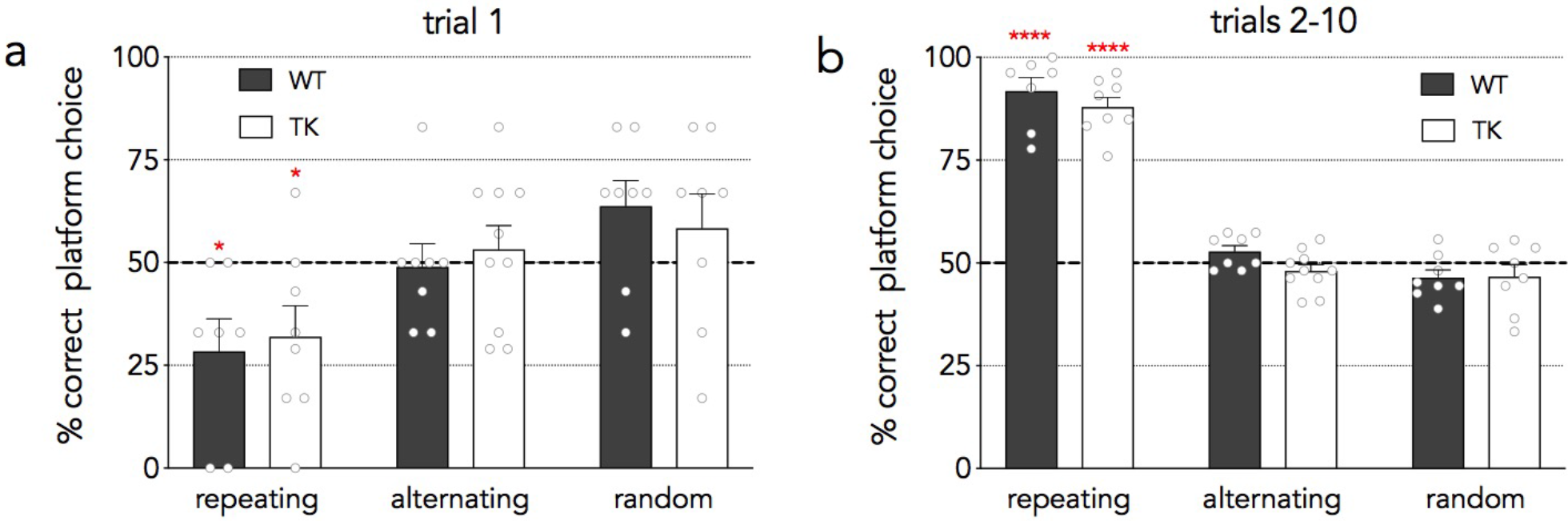
Correct platform choice. a) On trial 1, rats in the repeating condition swam to the correct location first at less than chance levels (50%), reflecting perseveration at the previous day’s location (one sample t-tests, *Ps<0.05). Rats in the alternating and random conditions swam to the correct platform first at chance levels (all Ps >0.05). In all conditions, WT and TK rats did not differ (Ps>0.9). b) On trials 2-10, only rats in the repeating condition swam to the correct location more than chance (****P<0.0001). WT and TK rats did not differ in any of the 3 conditions (all Ps>0.12).

### 3.5 Swim paths

The results thus far suggest that WT rats developed a more efficient but non-specific strategy to locate the escape platform in the alternating condition. To gain insight into the nature of the strategies employed by the rats, swim paths were inspected and are presented in Fig. 4 (additional examples in Supplementary Figs. 3-8). While rats in the repeating condition quickly learned to swim directly to the correct location, rats in the random and alternating conditions appeared to often approach the same platform location first on every trial, regardless of whether it was the correct or incorrect location. They would then typically check the other platform location and escape with variable success, depending on the accuracy of their trajectories.

**Figure 4:**
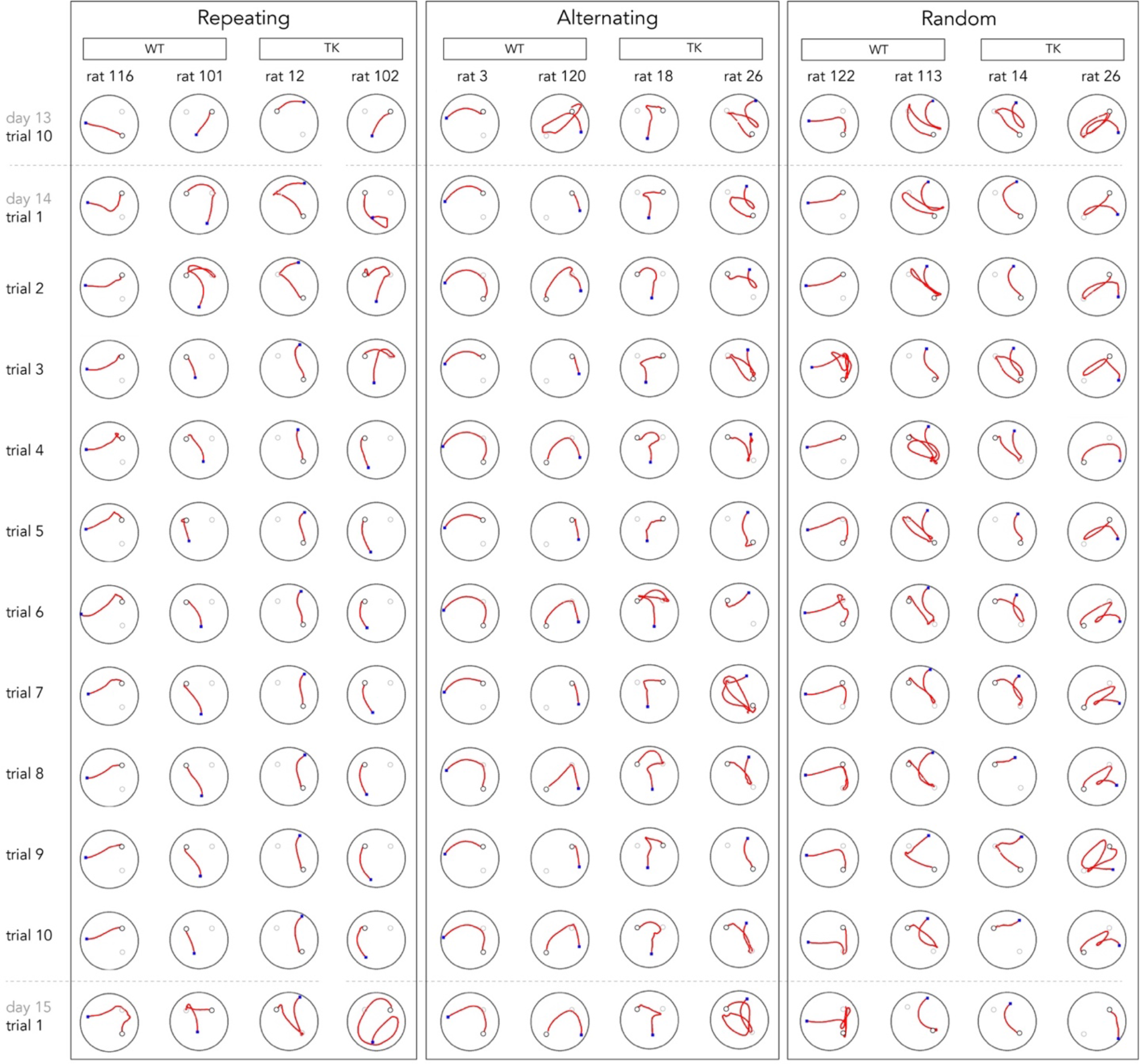
Swim paths on day 14. Rats in the repeating condition tended to swim directly to the platform after learning the platform location on a given day. WT rats in the alternating condition developed strong platform preferences and efficient escape strategies. TK rats in the alternating condition, and WT and TK rats in the random condition, developed weaker platform preferences and displayed less direct swim paths. Blue square indicates starting position for each trial; black circles indicate correct platform location; grey circles indicate incorrect/alternate platform location.

### 3.6 Navigation and spatial platform preference

To determine whether WT and TK rats differed in platform preference we quantified rats’ choice of the two locations irrespective of which one was reinforced on a given trial. Specifically, we calculated which platform location (40 cm zone) was approached first on a greater proportion of trials during days 10-15 (60 trials total/rat). The percentage of trials where the rats approached this platform first was used as a platform preference index. As expected, since rats in the repeating condition swam to the correct platform location on most trials, and the platform switched locations each day, they displayed only a small, but nonetheless significant, preference when trials were averaged over the 60 days (WT: 57%, TKs 54%; Fig. 5). In contrast, rats in the alternating condition developed a clear preference, which was greater in WT rats than in TK rats (87% vs 73%). The strong preference in WT rats emerged in the 2^nd^ half of training (Supplementary Fig. 9). Finally, rats in the random condition displayed a moderate platform preference that was not different between WT and TK rats (WT: 71%, TK: 73%). This analysis indicates that rats develop a habit-like preference for a specific platform location when escape options vary on a trial by trial basis. Furthermore, adult neurogenesis promotes this escape strategy when the location alternates perfectly.

**Figure 5:**
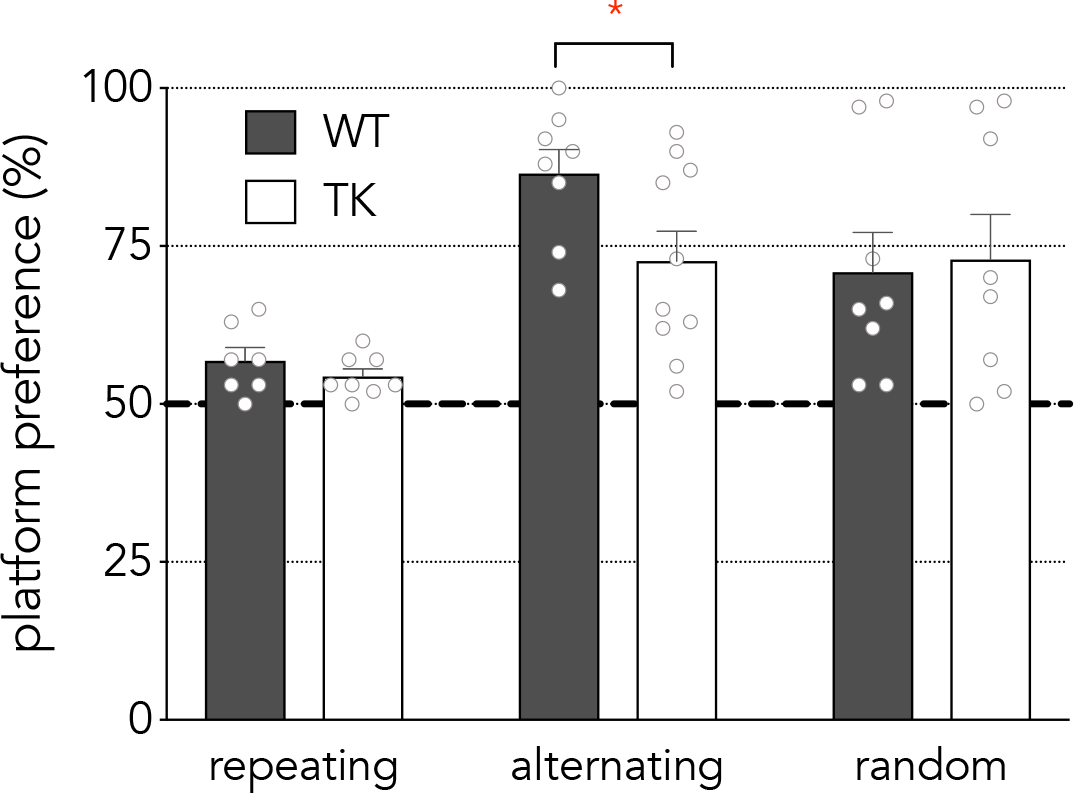
Percentage of trials where rats swam to their preferred platform location first. WT and TK rats in the repeating condition did not differ but swam to their preferred platform location on more than 50% of trials (WT vs TK: T_13_=1.1, P=0.3; both genotypes P < 0.05 vs chance). In the alternating condition, WT rats displayed a greater platform preference than TK rats (T_16_=2.2, *P=0.04). In the random condition, WT and TK rats developed a platform preference but did not differ from one another (T_14_=0.2, P=0.8).

To validate and extend the latency-based platform preference metric, we next analyzed the proximity to the preferred and non-preferred platform locations on days 10-15 (Fig. 6a-b). We focused on the alternating condition since this was the only condition where WT and TK rats differed in platform preference. We reasoned that platform biases should result in some trials where rats tend to swim closer to the correct platform location, and other trials where they swim closer to the incorrect location. Indeed, rats swam in closer proximity to the platform when it was in the preferred location as compared to the non-preferred location (“preferred” trials; Fig 6a). Proximity to the incorrect location showed a complementary pattern: rats avoided the incorrect area of the pool when the platform was in the preferred location as compared to the non-preferred location (Fig. 6b). However, this bias was weaker in TK rats since they swam in closer proximity to the incorrect location on preferred trials.

In addition, we examined heading angle relative to a direct path between the rat and the correct platform location, at 7 time points during the trial (Fig. 6c-d). On preferred trials, WT and TK rats performed similarly, and displayed a moderate heading angle error of 20-30° until they approached the platform at the end of the trial. On non-preferred trials, the heading angle error was much greater particularly in WT rats at the beginning of the trial. This is consistent with a strategy where WT rats first approach their preferred location and then, upon failing to find the platform, redirect their search to the non-preferred location. Blocking neurogenesis did not alter heading error in the repeating and random conditions (Supplementary Fig. 10).

**Figure 6:**
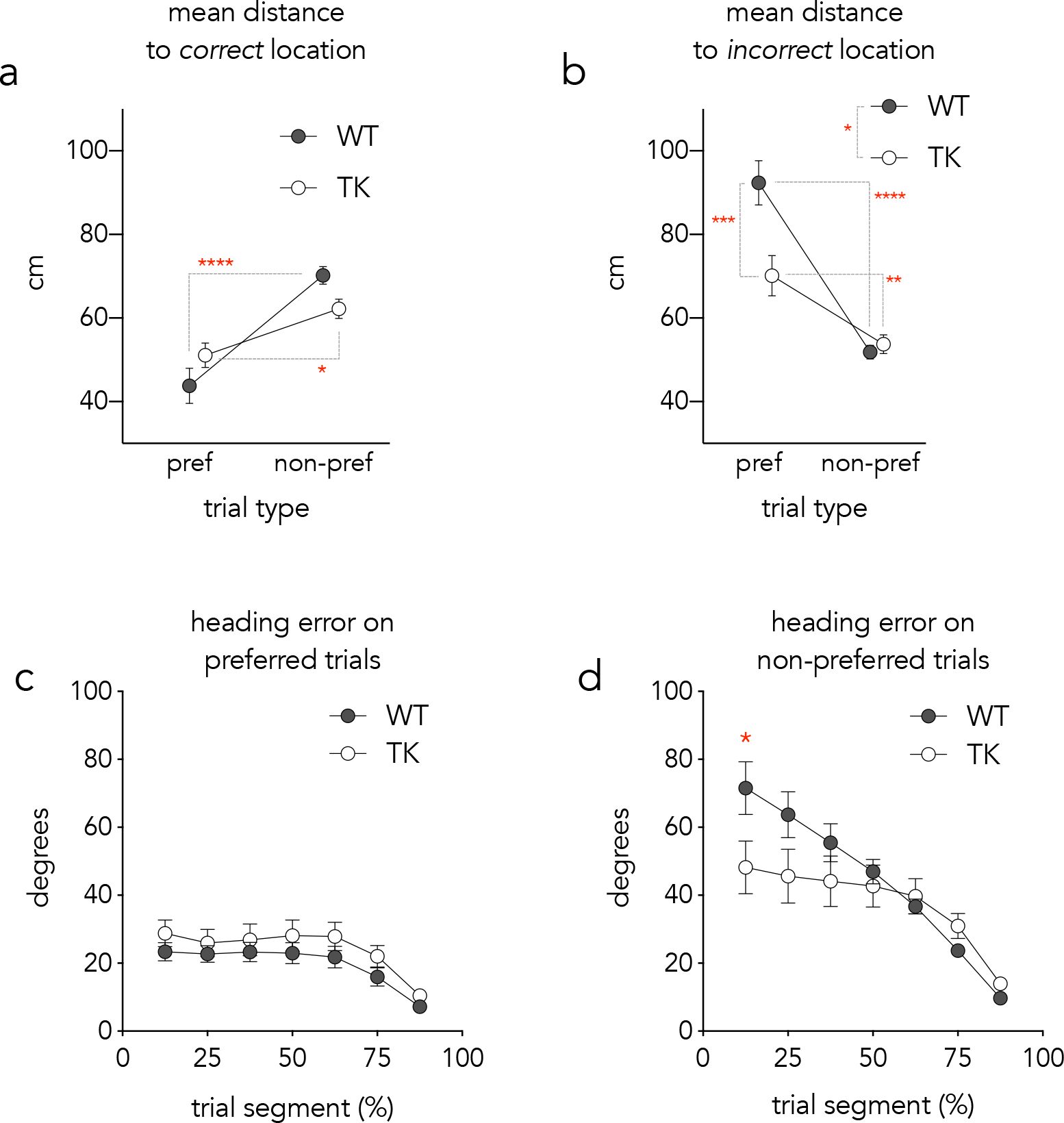
Navigation patterns as a function of platform location preference. a) Mean distance to the correct/actual platform location. WT and TK rats both swam in closer proximity to the platform on trials where it was in their preferred location compared to their non-preferred location (effect of preferred location F_1,16_=36, P<0.0001; genotype × preference interaction F_1,16_=6, P=0.03; post hoc comparison of preferred vs non-preferred location *P<0.05 and ****P<0.001). b) The mean distance from the incorrect/alternate platform location was greater for both WT and TK rats on trials where the platform was in the preferred location as compared to trials where it was located in the non-preferred location. WT rats’ distance from the incorrect platform was greater than TK rats, particularly on preferred-location trials (effect of preferred location F_1,16_=60, P<0.0001; effect of genotype F_1,16_=6, *P<0.05; genotype × preference interaction F_1,16_=11, P=0.005; **P<0.01, ****P<0.001). c) Heading error did not differ between WT and TK rats on trials where the platform was located in the rats’ preferred location (genotype effect F_1,16_=1.2, P=0.3). f) WT rats displayed greater initial heading error compared to TK rats on trials where the platform was in the non-preferred location (genotype × segment interaction, F_6,96_=5.7, P<0.0001; WT vs TK rats on trial 1, *P=0.03).

### 3.7 Search accuracy and efficiency

Having established that WT rats display a stronger platform preference in the alternating condition, we investigated how preference related to escape performance. With training, spatial search in the water maze progresses through a series of strategies that are increasingly spatially specific [5,23,24]. Here, rats were well-trained but the presence of a second, interfering escape location could have reduced spatial search specificity in TK rats. To test this we designed algorithms to differentiate direct, indirect and non-specific trajectories to the platform. In the alternating condition, we found that WT rats made more direct swims to the platform than TK rats, and this was entirely due to performance on preferred trials (Fig. 7a). This was also the case when we only analyzed preferred trials where rats approached the correct platform zone first, indicating that WT rats didn’t display more direct swims simply because they had a more consistent preference (T_16_=2.3, P=0.02). WT and TK rats made almost no direct swims when the platform was in the non-preferred location. Instead, rats tended to take indirect paths because they targeted their preferred location en route to the non-preferred location. Given their stronger platform preference, WT rats tended to make more indirect swims on nonpreferred platform trials (73% vs 58%, respectively; T_16_=1.7, P=0.1). There were no overall genotype differences in search strategies used in the repeating and random conditions (Supplementary Fig. 11). However, since the proximity analyses suggested that WT rats in the repeating condition perseverate less at the previous day’s platform location, we hypothesized that they would display more direct swims on trial 2. Indeed, WT rats swam directly to the platform on 57% of trials compared to 31% for TK rats (one tailed t-test, T_13_=1.9, P=0.04).

Since WT rats swam to their preferred platform more often and directly, we reasoned that they may also make fewer mistakes than TK rats on preferred trials. When the platform was in the preferred location WT rats approached both platform locations on only 15% of trials whereas TK rats approached both locations on 33% of preferred trials (40 cm zones; Fig. 6b). However, the genotype × trial type interaction was not significant. Additionally, we found no effect of platform preference on escape latencies or total 40 cm zone crossings/trial. Instead, WT rats were faster and made fewer errors on both preferred and non-preferred trials (Fig. 7c-d). These data collectively indicate that, while the neurogenesis-dependent platform preference enables optimal performance on the preferred trials, a generally superior spatial accuracy led to an overall enhanced performance of WT rats.

**Figure 7:**
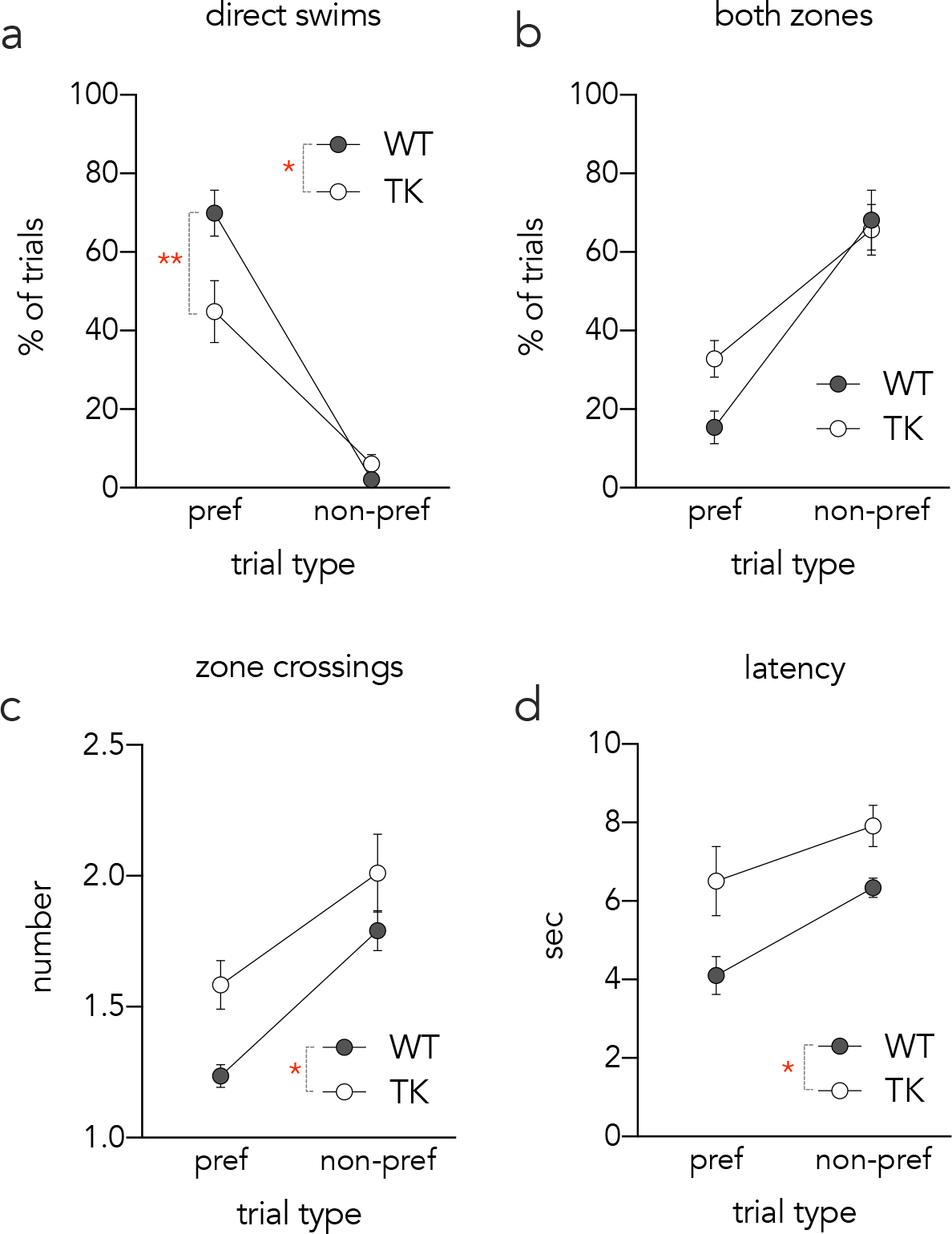
Escape path efficiency on preferred and non-preferred trials. a) Compared to TK rats, WT rats made more direct swims to the platform on preferred trials than non-preferred trials (genotype effect, F_1,16_=4.6, P=0.04; genotype × trial type interaction, F_1,16_=6.6, P=0.02; WT-preferred vs TK-preferred P=0.004). b) The proportion of trials where WT and TK rats visited both 40 cm platform location zones was not different (genotype effect, F_1,16_=2.6, P=0.12; genotype × trial type interaction, F_1,16_=2.1, P=0.17). c) TK rats made more 40 cm zone crossings per trial (genotype effect, F_1,16_=7.1, P=0.02; genotype × trial type interaction, F_1,16_=0.4, P=0.6). d) WT rats were faster to escape than TK rats, irrespective of whether the platform was is in the preferred or non-preferred location (genotype effect, F_1,16_=6.9, P=0.02; genotype × trial type interaction, F_1,16_=0.8, P=0.4). *P<0.05, **P<0.01.

## 4 Discussion

### 4.1 Summary of results

Here we investigated how adult neurogenesis regulates spatial search for goal locations that vary according to different spatiotemporal patterns. Given the proposed role of neurogenesis in mnemonic discrimination [27–29], and the hippocampal dependence of spatial delayed alternation [16], we hypothesized that blocking neurogenesis would prevent rats from learning an alternating pattern of platform locations, since there would be more interference between memories for recently-visited locations. We did not expect deficits in learning a repeating pattern, where the platform remains in a constant location within a day and therefore presents less interference, nor did we expect a role for neurogenesis in the random condition since there was no predictable pattern to be learned. These predictions were upheld in the sense that, aside from a transient reversal deficit in the repeating condition, blocking neurogenesis primarily impacted performance in the alternating condition. However, neurogenesis-intact WT rats did not escape faster due to superior alternation behavior since both genotypes alternated at chance levels. Instead, WT rats were generally more accurate in their search and less likely to approach and miss platform locations. Also, relative to TK rats, WT rats developed a more consistent strategy of swimming to the two possible escape locations in a fixed sequence, regardless of which location was reinforced. These search accuracy and strategy differences are unlikely to be due to motoric differences since swim speeds were similar and comparable results were obtained with multiple measures (latency, proximity, path length, heading angle, navigational strategy). Moreover, accuracy and strategy differences between WT and TK rats were not observed in the repeating and random conditions. These results therefore suggest that neurogenesis enables rats to develop optimal spatial search strategies when goal locations follow a spatiotemporal alternating pattern.

### 4.2 Adult neurogenesis promotes spatially precise search

Analyzing performance by platform location revealed no universal advantage of the sequential platform strategy in the alternating condition: WT rats were generally better on both preferred and non-preferred trials. Specifically, neurogenesis promoted faster escape by reducing the number of approaches (zone crossings) required to locate the platform. Whereas WT rats accurately targeted the platform, TK rats would often swim close to one platform zone only to change direction and then approach the other, suggesting less precise memory for the platform locations. Neurogenesis promoted more direct search paths on preferred-location trials, even after accounting for the weaker platform preference in TK rats. Neurogenesis-related accuracy on non-preferred trials was not detected by our strategy analyses. However, there was a non-significant tendency for TK rats to take fewer indirect paths and more nonspecific paths to the platform on non-preferred trials (suggesting that TK rats were less likely to sequentially target the preferred and non-preferred locations, and more likely to search randomly). While genotype differences were most pronounced in the alternating condition, in the repeating condition WT rats perseverated less at the previous day’s platform location and made more direct swims than TK rats on trial 2. Collectively, the enhanced spatial precision in WT rats is consistent with previous detailed analyses of search specificity in mice that showed that neurogenesis promotes spatially-specific search when learning a single escape location and when reversing to a new location [5]. Our data indicates that, even after many days of training, neurogenesis remains important for spatial search specificity when there are competing spatial goals.

Our findings are broadly consistent with a role for adult neurogenesis in the formation of precise and flexible hippocampal memories, which is typically revealed when animals are required to discriminate between similar, highly-interfering stimuli [27–29]. For example, loss and gain of function studies have identified a role for neurogenesis in contextual fear discrimination, where neurogenesis may promote context-specific fear by forming accurate memories for related environments [9–11,30]. Neurogenesis also enables rats to successfully learn conflicting lists of rewarded odors [12]. Finally, in spatial navigation paradigms, suppression of neurogenesis does not typically induce deficits in learning a single location [1–4]. However, when locations bear a high degree of similarity [31,32], or when animals are required to reverse a previously-learned association [5–8], neurogenesis-deficient animals are more likely to be impaired (but see [1,26]). Here, neurogenesis may have reduced interference between competing goals, and enabled more accurate search, by promoting detailed representations of spatial context and the platform locations. Why are TK rats less precise, even after 10+ days of training? This may be due to the powerful role that reinforcement plays in water maze learning: removing the platform on only 25% of trials more than doubles the escape latency and 50% reinforcement abolishes learning [33]. Possibly, less precise representations in TK rats, combined with reduced rates of reinforcement, led to a persistent inability to learn in the alternating condition. In contrast, in the repeating condition, consistently high rates of reinforcement at a single location may have enabled TK rats to overcome initial deficits. We note that alternating patterns present higher complexity and a greater working memory load, requiring more cognitive resources for successful encoding than repeating patterns [34]. In the random condition, unpredictable patterns of reinforcement may explain the relatively poor performance of both WT and TK rats. The differential performance between WT and TK rats in the alternating condition but not in the repeating or random conditions resembled Goldilocks effect [35], suggesting that neurogenesis benefits learning of not too simple or too complex, but moderately complex patterns.

### 4.3 Adult neurogenesis promotes a sequential search strategy

We hypothesized that neurogenesis would promote alternation learning, given that spatial alternation and nonmatch behaviors are hippocampal-dependent when there is a delay between trials [36,37]. Furthermore, previous reports have demonstrated that irradiated rats make more errors on a water maze-based delayed nonmatch to place task [18,38]. However, since these studies did not quantify spatial choice behavior, it is unclear if neurogenesis-intact rats performed better because they “nonmatched” or because they employed an efficient but nonspecific strategy as observed here. Our observations are consistent with a study showing that alternation is common in dry mazes but not water mazes [39], possibly because exploratory, win-shift behavior encourages discovery of new territory where rewards may be found. In contrast, escape from an aversive environment may benefit from a conservative strategy that reliably delivers the rat to a previous goal. Given the role of neurogenesis in modulating stress responses [40] and emotional behavior in ambiguous situations [41], differences in escape strategy choice could reflect neurogenesis functions in aversively-motivated behavior.

It is somewhat counterintuitive that neurogenesis promotes adoption of a fixed navigational trajectory, since the hippocampus is widely recognized for its role in promoting flexible behavior. The sequential platform strategy that is adopted by WT rats bears a strong resemblance to the inflexible, habit-like trajectories that are typically considered to be dependent upon the dorsolateral striatum rather than the hippocampus [13,14,42]. These response strategies are usually observed in dry mazes where rodents learn to perform simple navigational decisions (e.g. turn right, approach cue) over the course of many trials [42,43], consistent with the slow acquisition of a platform preference over ~10 days of training in alternating rats. From an adaptive perspective, this type of strategy is highly effective. Compared to an alternating strategy, the sequential platform strategy enables equally fast escape on 50% of trials and only slightly slower escape on the other 50% of trials. To the extent that it is “automated” and habit-like, it may substantially reduce cognitive load. Thus, it is plausible that neurogenesis facilitated control of behavior by the dorsolateral striatum, which led to an efficient, habit-like strategy in the alternating condition (see also evidence that the hippocampus can promote a response strategy [44]). To test this, future studies could pit place vs response strategies against each other by releasing animals from a novel location in the pool [45–47].

While the swim paths of WT rats in the alternating condition appeared to be inflexible and habit-like, an alternative explanation is that they reflected a systematic navigational plan. Numerous electrophysiological studies have revealed that hippocampal neurons encode sequences of spatial locations [15]. Both hippocampal pyramidal and granule neurons encode prospective information as rats plan future navigational behaviors [48–51]. Adult neurogenesis may therefore be involved in navigational planning, which is supported by recent evidence that new neurons promote choice of advantageous, but delayed, rewards. The swim paths would seem to support this idea since WT rats deployed smooth and reliable trajectories from the start point to the first platform and then the second platform. In contrast, TK rats often appeared to only decide once they had reached the midpoint of the two platform options, and they changed their course more frequently, resembling vicarious trial and error behaviors that reflect decision-making at choice points [13]. Similar patterns have been demonstrated in a human visual search task: whereas intact individuals display a systematic search strategy to reach a goal, hippocampal amnesics’ search is disorganized and inefficient [52].

## 5 Conclusions

Here we investigated the role of adult neurogenesis in navigation when goals shift according to various spatiotemporal patterns. We report two primary findings. First, adult neurogenesis was required for spatially precise search when goal locations alternated across days and when it alternated across trials, consistent with previous findings that neurogenesis promotes accurate memory and behavioral flexibility. Second, neurogenesis promoted a nonspecific but effective search strategy when the goal location alternated on each trial, where rats targeted goal locations sequentially in the same order on each trial. This novel finding suggests that new neurons promote efficient navigational strategies when there are competing goals, possibly by promoting habit-like responses or engaging hippocampal functions in future-oriented planning.

## Acknowledgements

The authors would like to thank Anna Schapiro for helpful discussion. This research was supported by a Discovery Grant from the Natural Sciences and Engineering Research Council of Canada (NSERC; JSS), a New Investigator Award from the Canadian Institutes of Health Research (JSS), a Scholar award from the Michael Smith Foundation for Health Research (JSS) and a NSERC graduate scholarship (RQY).

## References

[1] J.O. Groves, I. Leslie, G.-J. Huang, S.B. McHugh, A. Taylor, R. Mott, et al., Ablating Adult Neurogenesis in the Rat Has No Effect on Spatial Processing: Evidence from a Novel Pharmacogenetic Model, PLoS Genet. 9 (2013) e1003718. doi:10.1371/journal.pgen.1003718.s002.

[2] J.S. Snyder, N.S. Hong, R.J. McDonald, J.M. Wojtowicz, A role for adult neurogenesis in spatial long-term memory, Neuroscience. 130 (2005) 843–852. doi:10.1016/j.neuroscience.2004.10.009.

[3] T.M. Madsen, P.E.G. Kristjansen, T.G. Bolwig, G. Wörtwein, Arrested neuronal proliferation and impaired hippocampal function following fractionated brain irradiation in the adult rat, Neuroscience. 119 (2003) 635–642.

[4] T.J. Shors, D.A. Townsend, M. Zhao, Y. Kozorovitskiy, E. Gould, Neurogenesis may relate to some but not all types of hippocampal-dependent learning, Hippocampus. 12 (2002) 578–584. doi:10.1002/hipo.10103.

[5] A. Garthe, J. Behr, G. Kempermann, Adult-generated hippocampal neurons allow the flexible use of spatially precise learning strategies, PLoS ONE. 4 (2009) e5464. doi:10.1371/journal.pone.0005464.

[6] J.R. Epp, R. Silva Mera, S. Köhler, S.A. Josselyn, P.W. Frankland, Neurogenesis-mediated forgetting minimizes proactive interference, Nat Comms. 7 (2016) 10838. doi:10.1038/ncomms10838.

[7] N.S. Burghardt, E.H. Park, R. Hen, A.A. Fenton, Adult-born hippocampal neurons promote cognitive flexibility in mice, Hippocampus. 22 (2012) 1795–1808. doi:10.1002/hipo.22013.

[8] A.A. Swan, J.E. Clutton, P.K. Chary, S.G. Cook, G.G. Liu, M.R. Drew, Characterization of the role of adult neurogenesis in touch-screen discrimination learning, Hippocampus. 24 (2014) 1581–1591. doi:10.1002/hipo.22337.

[9] S. Tronel, L. Belnoue, N. Grosjean, J.-M. Revest, P.-V. Piazza, M. Koehl, et al., Adult-born neurons are necessary for extended contextual discrimination, Hippocampus. 22 (2012) 292–298. doi:10.1002/hipo.20895.

[10] A. Sahay, K.N. Scobie, A.S. Hill, C.M. O’Carroll, M.A. Kheirbek, N.S. Burghardt, et al., Increasing adult hippocampal neurogenesis is sufficient to improve pattern separation, Nature. 472 (2011) 466–470. doi:10.1038/nature09817.

[11] Y. Niibori, T.-S. Yu, J.R. Epp, K.G. Akers, S.A. Josselyn, P.W. Frankland, Suppression of adult neurogenesis impairs population coding of similar contexts in hippocampal CA3 region, Nat Comms. 3 (2012) 1253. doi:10.1038/ncomms2261.

[12] P. Luu, O.C. Sill, L. Gao, S. Becker, J.M. Wojtowicz, D.M. Smith, The role of adult hippocampal neurogenesis in reducing interference, Behav Neurosci. 126 (2012) 381–391. doi:10.1037/a0028252.

[13] A.D. Redish, Vicarious trial and error, Nat Rev Neurosci. 17 (2016) 147–159. doi:10.1038/nrn.2015.30.

[14] M.R. Penner, S.J.Y. Mizumori, Neural systems analysis of decision making during goal-directed navigation, Prog Neurobiol. 96 (2012) 96–135. doi:10.1016/j.pneurobio.2011.08.010.

[15] D.J. Foster, J.J. Knierim, Sequence learning and the role of the hippocampus in rodent navigation, Curr Opin Neurobiol. 22 (2012) 294–300. doi:10.1016/j.conb.2011.12.005.

[16] J.A. Ainge, M.A.A. van der Meer, R.F. Langston, E.R. Wood, Exploring the role of context-dependent hippocampal activity in spatial alternation behavior, Hippocampus. 17 (2007) 988–1002. doi:10.1002/hipo.20301.

[17] R.E. Hampson, L.E. Jarrard, S.A. Deadwyler, Effects of ibotenate hippocampal and extrahippocampal destruction on delayed-match and -nonmatch-to-sample behavior in rats, J Neurosci. 19 (1999) 1492–1507.

[18] G. Winocur, J.M. Wojtowicz, M. Sekeres, J.S. Snyder, S. Wang, Inhibition of neurogenesis interferes with hippocampus-dependent memory function, Hippocampus. 16 (2006) 296–304. doi:10.1002/hipo.20163.

[19] J.S. Snyder, L. Grigereit, A. Russo, D.R. Seib, M. Brewer, J. Pickel, et al., A Transgenic Rat for Specifically Inhibiting Adult Neurogenesis, eNeuro. 3 (2016). doi:10.1523/ENEURO.0064-16.2016.

[20] R. Morris, Developments of a water-maze procedure for studying spatial learning in the rat, J Neurosci Methods. 11 (1984) 47–60.

[21] M. Gallagher, R. Burwell, M. Burchinal, Severity of spatial learning impairment in aging: development of a learning index for performance in the Morris water maze, Behav Neurosci. 107 (1993) 618–626.

[22] D.A. Hamilton, C.S. Rosenfelt, I.Q. Whishaw, Sequential control of navigation by locale and taxon cues in the Morris water task, Behav Brain Res. 154 (2004) 385–397. doi:10.1016/j.bbr.2004.03.005.

[23] S. Ruediger, D. Spirig, F. Donato, P. Caroni, Goal-oriented searching mediated by ventral hippocampus early in trial-and-error learning, Nat Neurosci. 15 (2012) 1563–1571. doi:10.1038/nn.3224.

[24] J. Gil-Mohapel, P.S. Brocardo, W. Choquette, R. Gothard, J.M. Simpson, B.R. Christie, Hippocampal neurogenesis levels predict WATERMAZE search strategies in the aging brain, PLoS ONE. 8 (2013) e75125. doi:10.1371/journal.pone.0075125.

[25] I. Tomás Pereira, R.D. Burwell, Using the spatial learning index to evaluate performance on the water maze, Behav Neurosci. 129 (2015) 533–539. doi:10.1037/bne0000078.

[26] D.R. Seib, E. Chahley, O. Princz-Lebel, J.S. Snyder, Intact memory for local and distal cues in male and female rats that lack adult neurogenesis, PLoS ONE. 13 (2018) e0197869–15. doi:10.1371/journal.pone.0197869.

[27] J.B. Aimone, W. Deng, F.H. Gage, Resolving new memories: a critical look at the dentate gyrus, adult neurogenesis, and pattern separation, Neuron. 70 (2011) 589–596. doi:10.1016/j.neuron.2011.05.010.

[28] S. Becker, Neurogenesis and pattern separation: time for a divorce, WIREs Cogn Sci. 8 (2017) e1427. doi:10.1002/wcs.1427.

[29] A. Sahay, D.A. Wilson, R. Hen, Pattern separation: a common function for new neurons in hippocampus and olfactory bulb, Neuron. 70 (2011) 582–588. doi:10.1016/j.neuron.2011.05.012.

[30] M.A. Kheirbek, L. Tannenholz, R. Hen, NR2B-dependent plasticity of adult-born granule cells is necessary for context discrimination, J Neurosci. 32 (2012) 8696–8702. doi:10.1523/JNEUROSCI.1692-12.2012.

[31] C.D. Clelland, M. Choi, C. Romberg, G.D. Clemenson, A. Fragniere, P. Tyers, et al., A functional role for adult hippocampal neurogenesis in spatial pattern separation, 325 (2009) 210–213. doi:10.1126/science.1173215.

[32] P. Bekinschtein, B.A. Kent, C.A. Oomen, G.D. Clemenson, F.H. Gage, L.M. Saksida, et al., Brain-derived neurotrophic factor interacts with adult-born immature cells in the dentate gyrus during consolidation of overlapping memories, Hippocampus. 24 (2014) 905–911. doi:10.1002/hipo.22304.

[33] C.L. Gonzalez, B. Kolb, I.Q. Whishaw, A cautionary note regarding drug and brain lesion studies that use swimming pool tasks: partial reinforcement impairs acquisition of place learning in a swimming pool but not on dry land, Behav Brain Res. 112 (2000) 43–52.

[34] R.Q. Yu, D. Osherson, J. Zhao, Alternation blindness in the representation of binary sequences, J Exp Psychol Hum Percept Perform. 44 (2018) 493–502. doi:10.1037/xhp0000476.

[35] C. Kidd, S.T. Piantadosi, R.N. Aslin, The Goldilocks effect: human infants allocate attention to visual sequences that are neither too simple nor too complex, PLoS ONE. 7 (2012) e36399. doi:10.1371/journal.pone.0036399.

[36] J.T.B.G.E.H. David S Olton, Hippocampus, space, and memory, Behav Brain Sci. 2 (1979) 313–365.

[37] P.A. Dudchenko, An overview of the tasks used to test working memory in rodents, Neuroscience and Biobehavioral Reviews. 28 (2004) 699–709. doi:10.1016/j.neubiorev.2004.09.002.

[38] G. Winocur, J.M. Wojtowicz, J. Huang, I.F. Tannock, Physical exercise prevents suppression of hippocampal neurogenesis and reduces cognitive impairment in chemotherapy-treated rats, Psychopharmacology (Berl). 231 (2013) 2311–2320. doi:10.1007/s00213-013-3394-0.

[39] I.Q. Whishaw, T.J. Pasztor, Rats alternate on a dry-land but not swimming-pool (Morris task) place task: implications for spatial processing, Behav Neurosci. 114 (2000) 442–446.

[40] J.S. Snyder, A. Soumier, M. Brewer, J. Pickel, H.A. Cameron, Adult hippocampal neurogenesis buffers stress responses and depressive behaviour, Nature. 476 (2011) 458–461. doi:10.1038/nature10287.

[41] L.R. Glover, T.J. Schoenfeld, R.-M. Karlsson, D.M. Bannerman, H.A. Cameron, Ongoing neurogenesis in the adult dentate gyrus mediates behavioral responses to ambiguous threat cues, PLoS Biol. 15 (2017) e2001154. doi:10.1371/journal.pbio.2001154.

[42] M.G. Packard, J.L. McGaugh, Inactivation of hippocampus or caudate nucleus with lidocaine differentially affects expression of place and response learning, Neurobiol Learn Mem. 65 (1996) 65–72. doi:10.1006/nlme.1996.0007.

[43] Q. Chang, P.E. Gold, Switching memory systems during learning: changes in patterns of brain acetylcholine release in the hippocampus and striatum in rats, J Neurosci. 23 (2003) 3001–3005.

[44] J. Ferbinteanu, Contributions of Hippocampus and Striatum to Memory-Guided Behavior Depend on Past Experience, J Neurosci. 36 (2016) 6459–6470. doi:10.1523/JNEUROSCI.0840-16.2016.

[45] H. Eichenbaum, C. Stewart, R.G. Morris, Hippocampal representation in place learning, J Neurosci. 10 (1990) 3531–3542.

[46] R.J. McDonald, N.M. White, Parallel information processing in the water maze: evidence for independent memory systems involving dorsal striatum and hippocampus, Behav Neural Biol. 61 (1994) 260–270.

[47] J.S. Snyder, S.P. Cahill, P.W. Frankland, Running promotes spatial bias independently of adult neurogenesis, Hippocampus. 31 (2017) 15113. doi:10.1002/hipo.22737.

[48] Dentate network activity is necessary for spatial working memory by supporting CA3 sharp-wave ripple generation and prospective firing of CA3 neurons, Nat Neurosci. 33 (2018) 1. doi:10.1038/s41593-017-0061-5.

[49] B.E. Pfeiffer, D.J. Foster, Hippocampal place-cell sequences depict future paths to remembered goals, Nature. 497 (2013) 74–79. doi:10.1038/nature12112.

[50] A. Johnson, A.D. Redish, Neural ensembles in CA3 transiently encode paths forward of the animal at a decision point, J Neurosci. 27 (2007) 12176–12189. doi:10.1523/JNEUROSCI.3761-07.2007.

[51] J. Ferbinteanu, M.L. Shapiro, Prospective and retrospective memory coding in the hippocampus, Neuron. 40 (2003) 1227–1239.

[52] L.T.S. Yee, D.E. Warren, J.L. Voss, M.C. Duff, D. Tranel, N.J. Cohen, The hippocampus uses information just encountered to guide efficient ongoing behavior, Hippocampus. 24 (2014) 154–164. doi:10.1002/hipo.22211.

