## Supplementary material for "Adult neurogenesis promotes efficient, nonspecific search strategies in a spatial alternation water maze task"

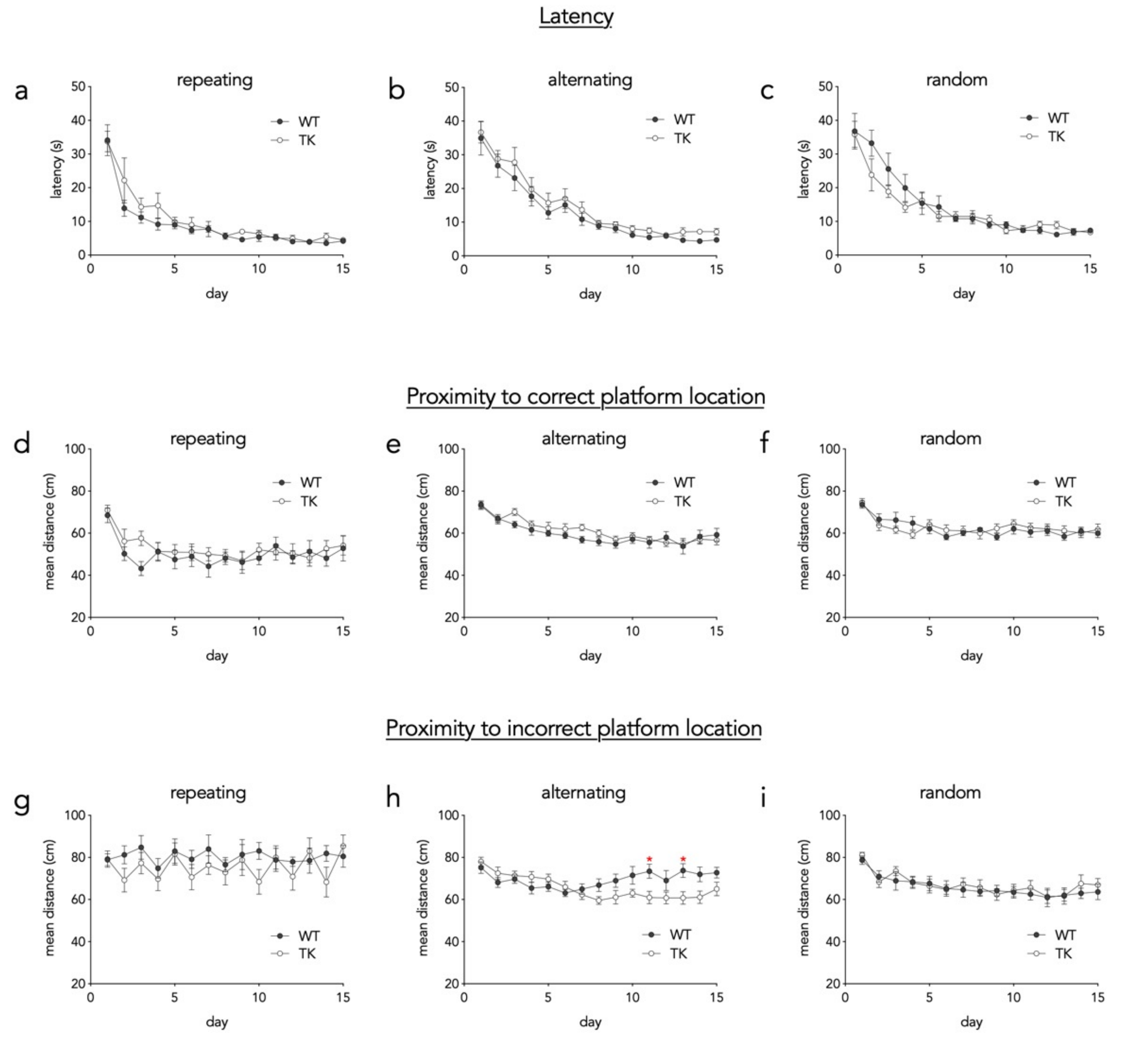


Supplementary Figure 1: Task acquisition. A-C) Latency to reach the platform over the full 15 days of training. For all conditions, escape latencies decreased over days and were not different between genotypes (effect of day, all Ps <0.0001; effect of genotype, all Ps > 0.17). D-F) Proximity to the correct platform location decreased over the 15 days of training and was not different between WT and TK rats (effect of day, all Ps <0.0001; effect of genotype, all Ps > 0.3). G-I) Proximity to the incorrect platform location over the full 15 days of training. In the repeating condition there was no difference across days or genotypes (both Ps > 0.2). In the alternating condition there was an effect of day (P < 0.001), no effect of genotype (P = 0.2) and a day x genotype interaction (P < 0.0001). TK rats' average distance from the incorrect platform was less than WT rats on days 11 and 13 (*P < 0.05). In the random condition there was an effect of day (P < 0.0001) but not genotype (P = 0.7).

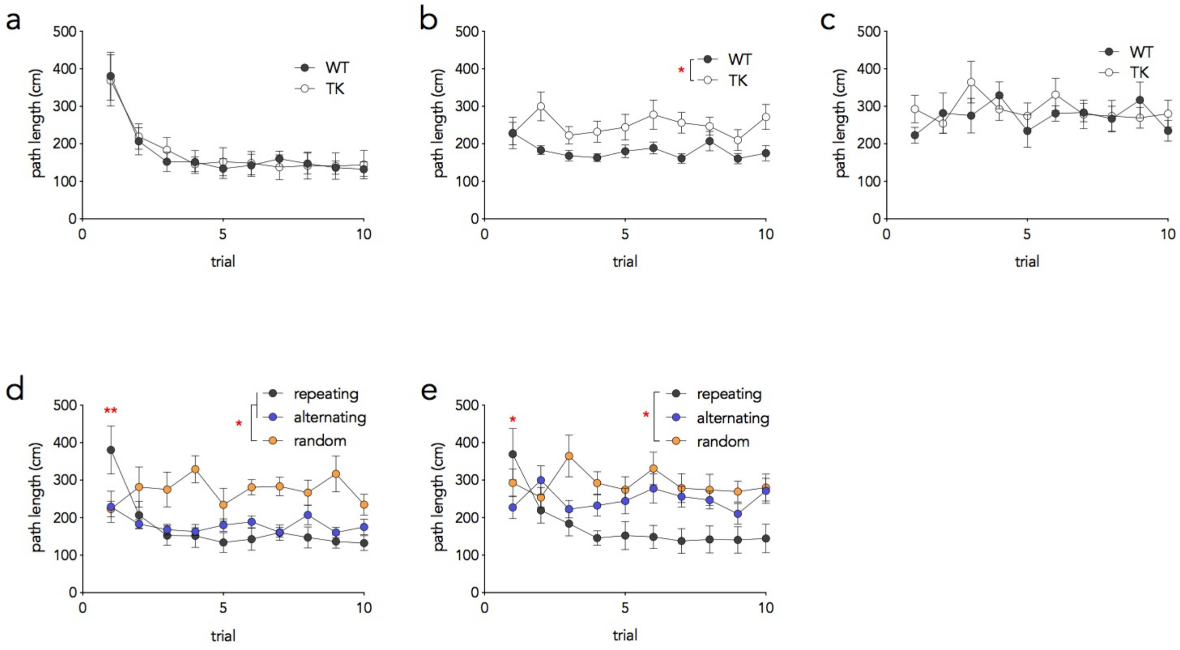


<u>Supplementary Figure 2: Trial path lengths on days 10-15</u>. a) WT and TK rats in the repeating condition did not differ, and required 1-2 trials to reach asymptotic performance (effect of genotype, $F_{1,14}$=0.01, P=0.9; effect of trial, $F_{9,117}$=23). b) In the alternating condition, WT rats swam shorter distances to reach the platform (effect of genotype, $F_{1,16}$=6.3, *P=0.02). c) In the random condition, WT and TK rats swam equal distances to reach the platform ($F_{1,14}$=0.2, P=0.6). d) WT rats in the random condition swam greater distances to locate the platform that rats in the repeating and alternating conditions (effect of condition, $F_{2,21}$=6.3, P=0.008; *P<0.05 vs alternating and repeating groups). On trial 1, however, rats in the repeating condition swam longer paths than both alternating and random groups (condition x trial interaction, $F_{18,189}$=4.9, P<0.0001; post hoc comparisons both **P<0.01). e) TK rats in the repeating condition generally swam farther to locate the platform than rats in the random condition (effect of condition, $F_{2,23}$=4.1, P=0.03; repeating vs random *P=0.02, repeating vs alternating P=0.17). On trial 1, TK rats in the repeating condition swam farther to locate the platform than rats in the alternating condition (condition x trial interaction, $F_{18,207}$=4.1, P<0.0001; post hoc comparison *P<0.05).

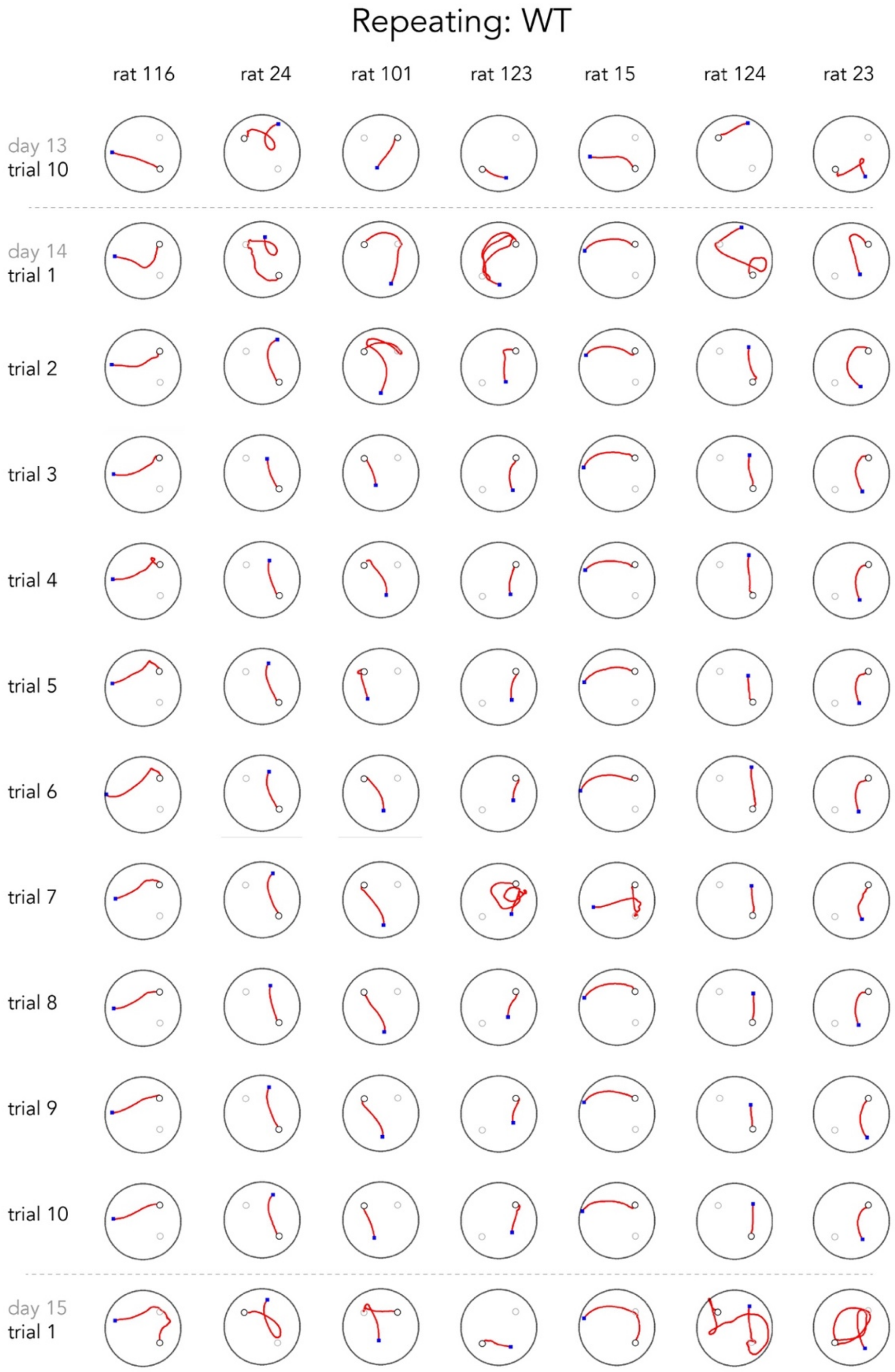


Supplementary Figure 3: Swim paths for WT rats in the repeating condition.

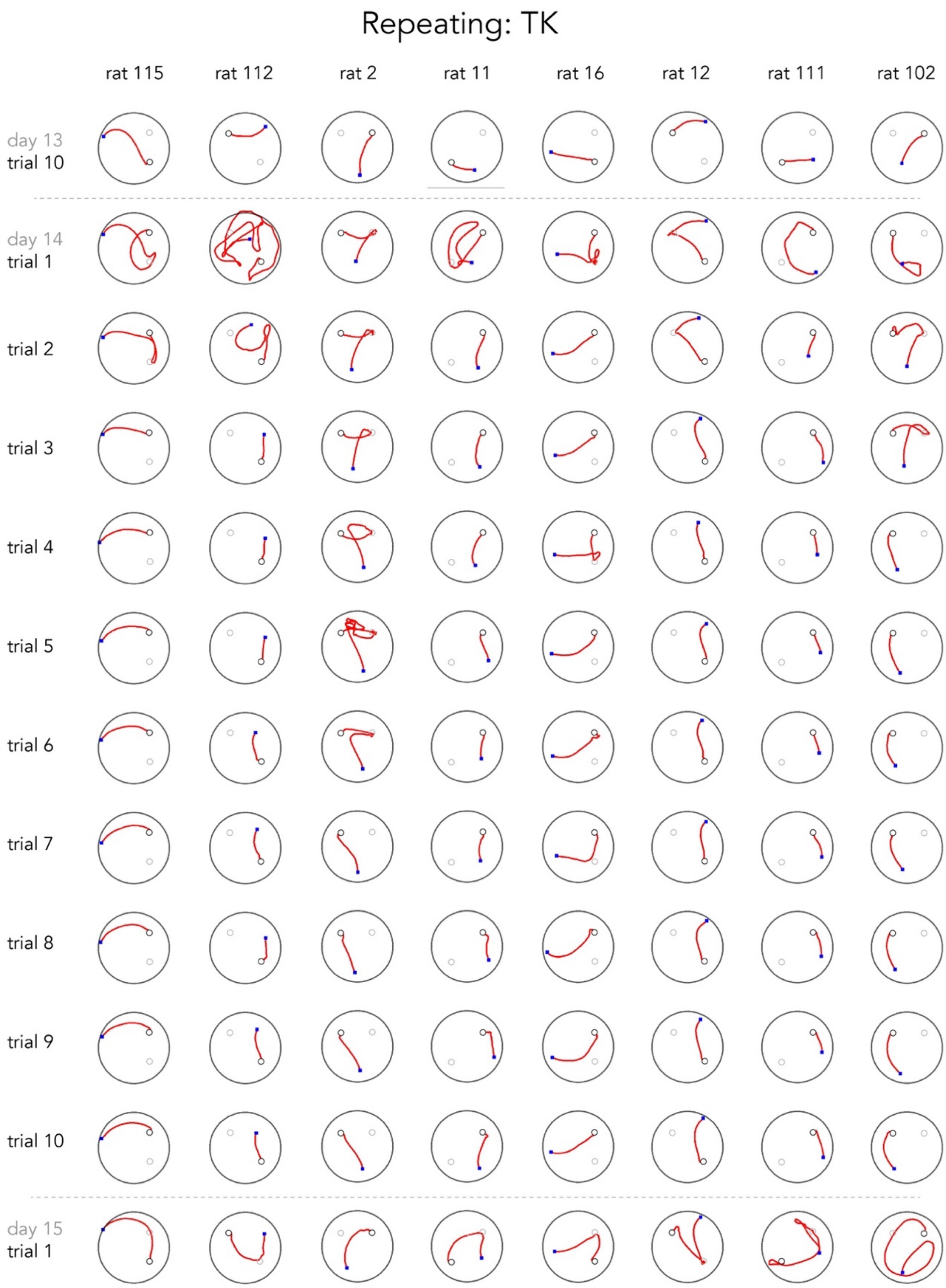


Supplementary Figure 4: Swim paths for TK rats in the repeating condition.

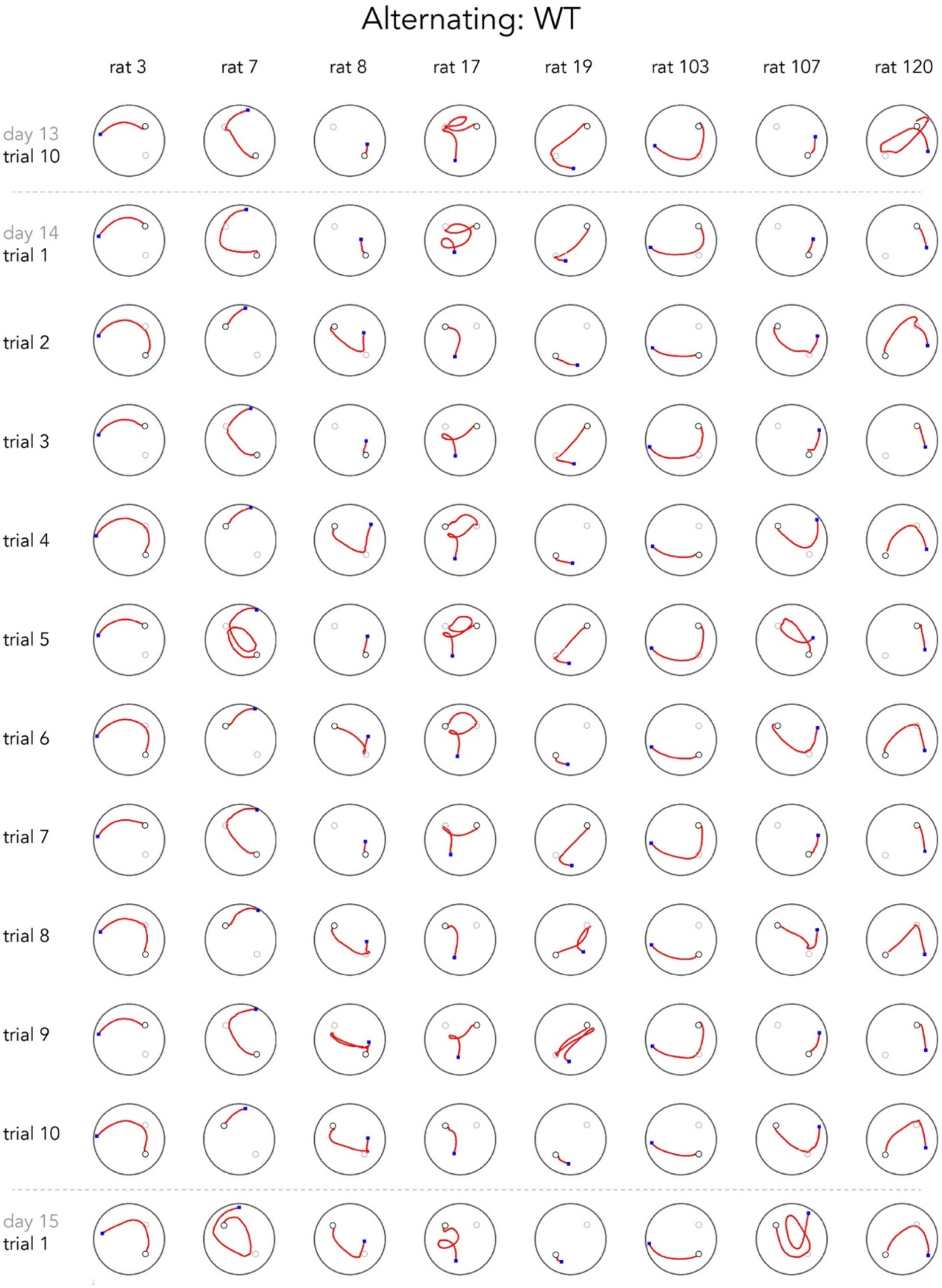


Supplementary Figure 5: Swim paths for WT rats in the alternating condition.

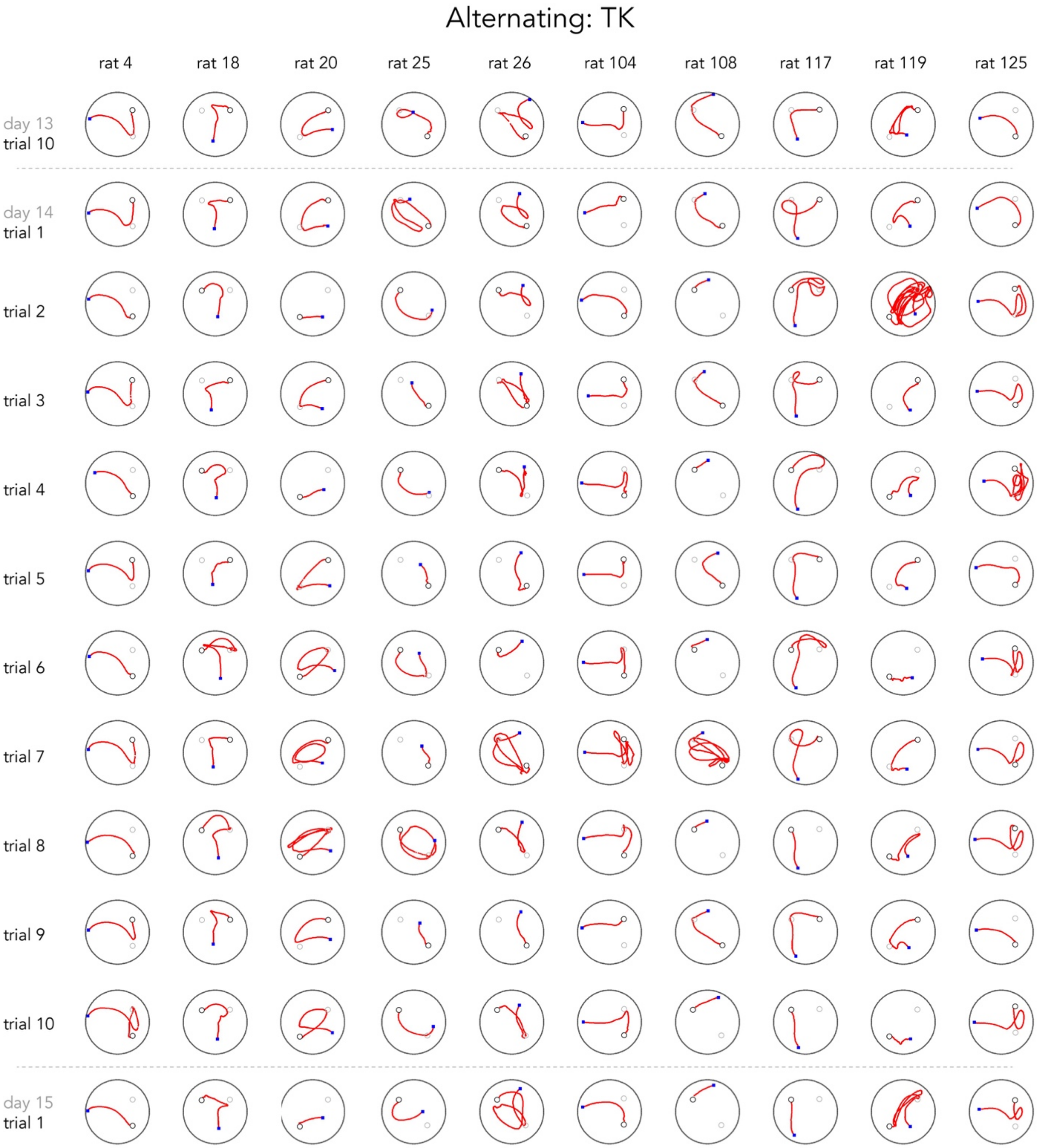


Supplementary Figure 6: Swim paths for TK rats in the alternating condition.

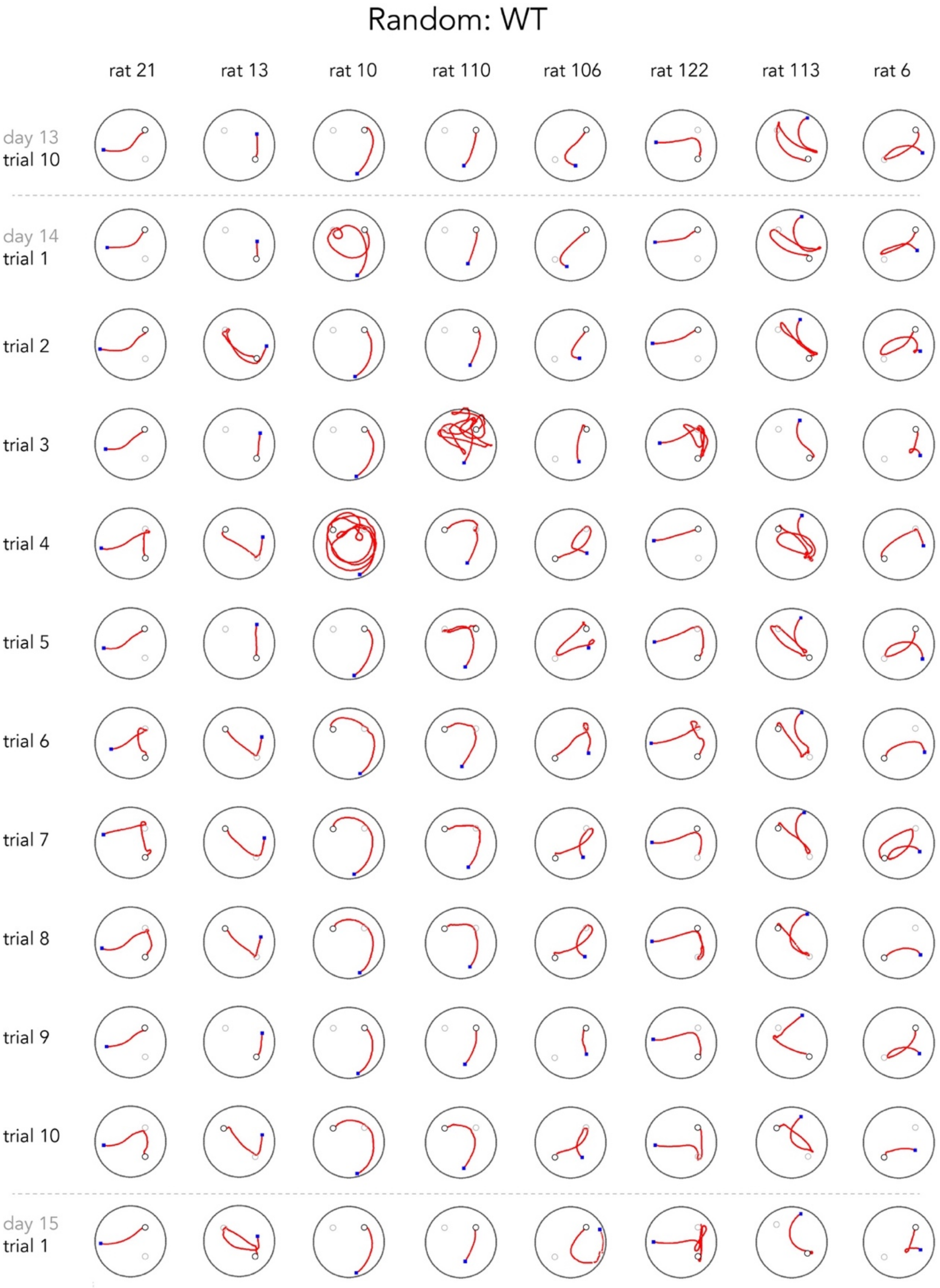


Supplementary Figure 7: Swim paths for WT rats in the random condition.

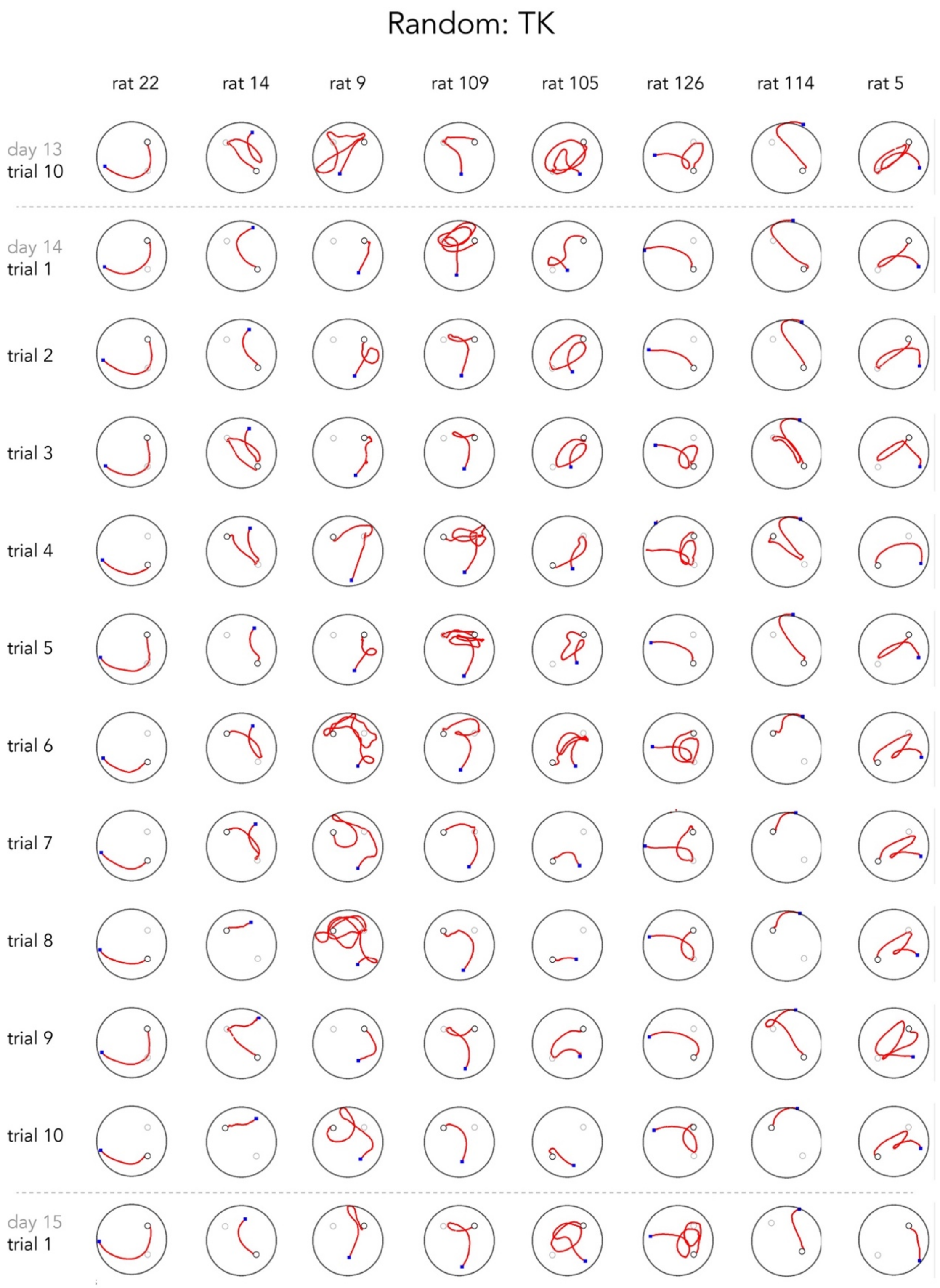


Supplementary Figure 8: Swim paths for TK rats in the random condition.

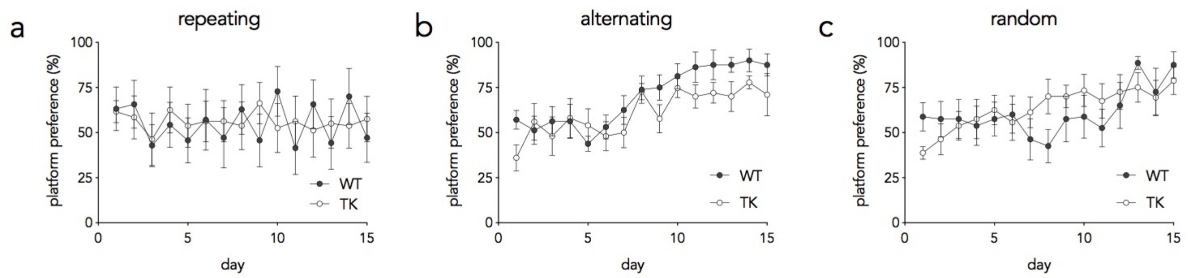


Supplementary Figure 9: Platform preference over the full 15 days of testing. a) In the repeating condition, platform preference scores were not different over days or across genotypes (day effect, $F_{14,182}$=0.4, P=0.98; genotype effect, $F_{1,13}$=0.06, P=0.8, interaction $F_{14,182}$=0.4, P=0.97). b) In the alternating condition, preference scores increased across days of training in WT and TK rats (day effect, $F_{14,224}$=6.6, P<0.0001; genotype effect, $F_{1,16}$=4.2, P=0.06). c) In the random condition, preference scores increased across days of training in WT and TK rats (day effect, $F_{14,96}$=3.6, P<0.0001; genotype effect, $F_{1,14}$=0.08, P=0.8).

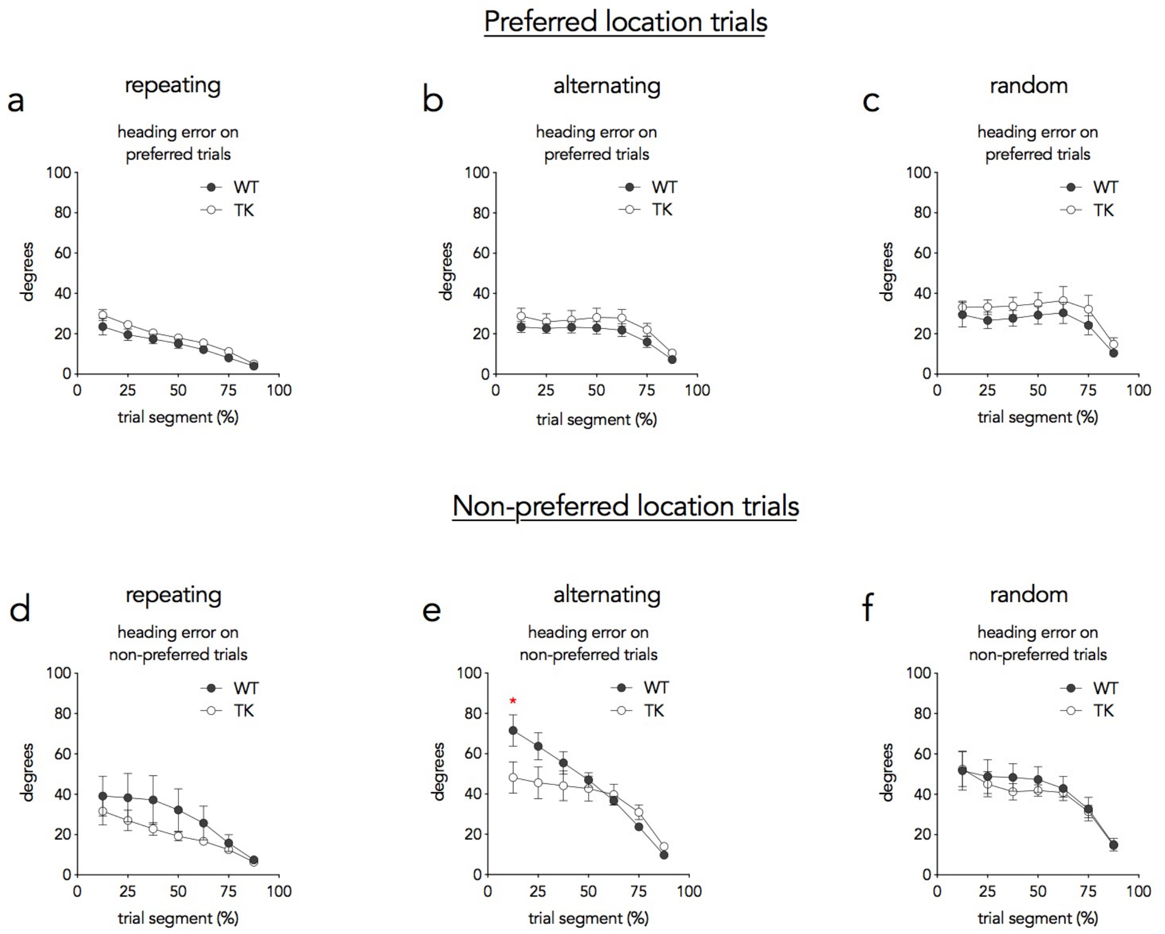


Supplementary Figure 10: Heading error for trials where the platform was located in the preferred and non-preferred locations. While heading error consistently decreased across trial segment, there were no genotype differences or genotype x segment interactions (all Ps>0.15), except for the alternating condition when the platform was in the non-preferred location (e; see main text).

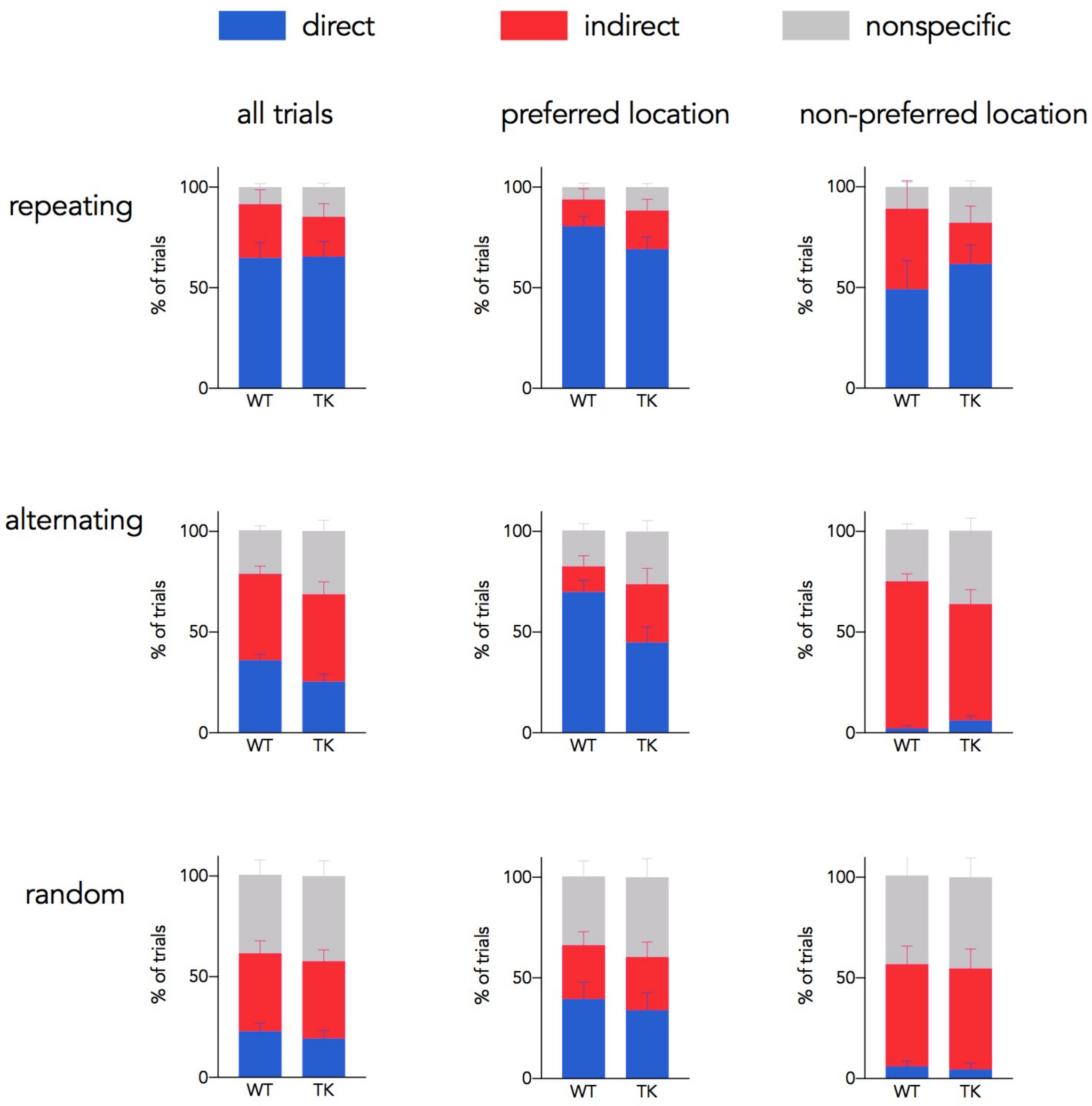


Supplementary Figure 11: Search strategy breakdown by genotype, condition, and trial type.
